# On the Reconstruction of Magnetic Resonance Current Density Images of the Human Brain: Pitfalls and Perspectives

**DOI:** 10.1101/2021.04.22.440915

**Authors:** Hasan H. Eroğlu, Oula Puonti, Cihan Göksu, Fróði Gregersen, Hartwig R. Siebner, Lars G. Hanson, Axel Thielscher

## Abstract

Magnetic resonance current density imaging (MRCDI) of the human brain aims to reconstruct the current density distribution caused by transcranial electric stimulation from MR-based measurements of the current-induced magnetic fields. The reconstruction problem is challenging due to a low signal-to-noise ratio and a limited volume coverage of the MR-based measurements, the lack of data from the scalp and skull regions and because MRCDI is only sensitive to the component of the current-induced magnetic field parallel to the scanner field. Most existing reconstruction approaches have been validated using simulation studies and measurements in phantoms with simplified geometries. Only one reconstruction method, the projected current density algorithm, has been applied to human in-vivo data so far, however resulting in blurred current density estimates even when applied to noise-free simulated data.

We analyze the underlying causes for the limited performance of the projected current density algorithm when applied to human brain data. In addition, we compare it with an approach that relies on the optimization of the conductivities of a small number of tissue compartments of anatomically detailed head models reconstructed from structural MR data. Both for simulated ground truth data and human in-vivo MRCDI data, our results indicate that the estimation of current densities benefits more from using a personalized volume conductor model than from applying the projected current density algorithm. In particular, we introduce a hierarchical statistical testing approach as a principled way to test and compare the quality of reconstructed current density images that accounts for the limited signal-to-noise ratio of the human in-vivo MRCDI data and the fact that the ground truth of the current density is unknown for measured data. Our results indicate that the statistical testing approach constitutes a valuable framework for the further development of accurate volume conductor models of the head. Our findings also highlight the importance of tailoring the reconstruction approaches to the quality and specific properties of the available data.

## INTRODUCTION

Knowledge of electrical current density (***J***) and conductivity (σ) distributions in the human brain is important in many neuroscience applications. It enables the control and optimization of the brain areas targeted by transcranial brain stimulation, and is needed for localizing neural sources from electro- and magnetoencephalography data. Also, it might be useful for characterizing malignant tissue (Holdefer et al., 2006; Miranda et al., 2014; Nathan et al., 1993). Magnetic resonance current density imaging (MRCDI) and magnetic resonance electrical impedance tomography (MREIT) are two modalities using transcranial current injections and MR imaging (MRI) to reconstruct current density and conductivity distributions in the brain (Scott et al., 1991). In both methods, the currents are injected in synchrony with an MRI pulse sequence to create a magnetic flux density distribution that changes the magnetic field of the MR scanner. As a result, the phase of the measured MR signal is modulated by the z-component of the current-induced magnetic flux density (*B_z_*), i.e. the part that is parallel to the static scanner field. The MR phase information thus can be used to obtain cross-sectional images of the current-induced magnetic flux density *B_z_*.

Several methods have been developed to reconstruct the current density distribution from the measured *B_z_* images. Considering Ampere’s law, a unique and unambiguous reconstruction requires knowledge of all three components of the current-induced magnetic flux density. As only the *B_z_* component parallel to the main magnetic field of the MRI scanner can be measured during an MRCDI experiment, accurate and complete current density mapping requires rotating the measured object inside the scanner (Eyüboğlu, 2006; Scott et al., 1991; Woo and Seo, 2008), which is impractical for human in-vivo brain imaging. Alternative methods thus aim to reconstruct the two components of the electrical current density that are orthogonal to the main scanner field (i.e., *J_x_* and *J_y_*) from a single *B_z_* image (Ider et al., 2010; Jeong et al., 2014; Park et al., 2007). In particular, the “projected current density” method introduced by Jeong et al (2014) was used in recent studies on in-vivo human brain MRCDI (Chauhan et al., 2018; Göksu et al., 2018a; Kasinadhuni et al., 2017). Combined with data from diffusion tensor imaging, projected current densities estimated with this method for two current injection directions (left-right and anterior-posterior) have also been used to reconstruct the ohmic conductivity distribution in the human brain (Chauhan et al., 2018). The accuracy of the reconstructed current density distribution depends on both the quality of the *B_z_* images and the properties of the reconstruction methods. Usually, their performances are good for simple phantoms with one or more conductivity inhomogeneities inside a homogenous well-conductive background (Kwon et al., 2007; Oh et al., 2003; Park et al., 2007). Also the reconstruction method used so far for in-vivo human brain MRCDI studies was validated in this way (Jeong et al., 2014). However, this situation is strongly simplified compared to MRCDI measurements of the human head, which has a complex anatomy and where the brain is surrounded by the highly resistive skull and conductive scalp.

In an earlier study (Göksu et al., 2018), we observed that the reconstructed current densities were only coarse estimates of the current flow in the brain even when applied to noise-free simulated data. Here, we explore the underlying causes and reveal fundamental limitations that occur when applying the projected current density method of Jeong et al (2014) to *B_z_* image of the human brain. We then explore whether the method can be modified to achieve better reconstruction performance. We also test an alternative approach to estimate the current density distribution that is based on the optimization of the conductivities of an anatomically detailed head model to fit the measured *B_z_* image. Using both simulated *B_z_* data serving as ground truth and *B_z_* data from human in-vivo measurements, we show that the alternative approach performs better than the projected current density method. In particular, we advocate for systematic statistical evaluations of the goodness of fit between measured and simulated *B_z_* data as a principled way to draw conclusions on the accuracy of simulations. A preprint of this paper was published on biorxiv.

## MATERIAL AND METHODS

### Human Brain Imaging

Two healthy volunteers, who had no previous neurological and psychiatric disorders, were included in this study. Written informed consent was obtained from the participants prior to the scans and they were screened for contraindications to MRI and transcranial electric stimulation (TES). The study complied with the Helsinki declaration on human experimentation and was approved by the Ethics Committee of the Capital Region of Denmark. We injected electrical current with ±1 mA magnitude in the left-right (LR) and anterior-posterior (AP) directions via surface electrodes attached to the head. The currents were applied in synchrony with a multigradient-echo MR pulse sequence (Göksu et al., 2018a; Göksu et al., 2018b) on a 3T MRI scanner equipped with a 64-channel head coil (Magnetom PRISMA, Siemens). We measured *B_z_* distributions by post processing the acquired MR phase images (Göksu et al., 2018a) and corrected unwanted effects of the magnetic stray field caused by the cable currents on the *B_z_* measurements (Göksu et al., 2019). The *B_z_* measurements were denoised with a Gaussian filter with a full width at half maximum (FWHM) value of 3 pixels.

We generated 3D individualized head models of the subjects by utilizing prior structural T1- and T2-weighted measurements (Nielsen et al., 2018) to simulate the current density distribution in the head and the resulting magnetic field distribution caused by the currents. As described in more detail below, we used different head models of increasing anatomical complexity (single compartment, three compartments, four compartments) for the simulations.

### Forward Simulations

Current densities were calculated using the Finite Element Method (FEM) implemented in SimNIBS 3.1.0 (Thielscher et al., 2015). The initial part of the study relied on the head model *ernie* included in the example dataset. This volume conductor model has a high resolution and is composed of 4.58×10^6^ tetrahedral elements with an average volume of 1 mm^3^. For the later part, individual head models with similarly high resolutions were reconstructed from T1-weighted and T2-weighted structural MR images. Detailed explanations on the construction of the head model are given in (Nielsen et al., 2018).

For the full models with all tissue types, the ohmic conductivities were assigned as σ*_scalp_* = 0.465, σ*^skull^* = 0.01, σ*_WM_* = 0.126, σ*_GM_* = 0.275, and σ*_CSF_* = 1.654 S/m (Fig. 1A) (Thielscher et al., 2011). In addition, we simulated anatomically simplified models by combining tissue types. A single compartment (1c, i.e. homogenous) model was used by setting all conductivities to σ*_o_* = 1 S/m. A model with three compartments (3c) was created by assigning the same conductivity σ*_inside_* = 0.34 S/m to CSF, GM and WM (σ*_scalp_* = 0.465 S/m, σ*_skull_* = 0.01 S/m). The conductivity σ*_inside_* was calculated by taking the average conductivity of the region composed of WM, GM, and CSF volumes, weighted according to their relative volumes, in the slice being imaged. Finally, a model with four compartments (4c) was created by combining GM and WM with a conductivity of σ*_brain_* = 0.18 S/m that was determined as the average of σ*_WM_* and σ*_GM_* in the imaged slice (σ*_scalp_* = 0.465, σ*_skull_* = 0.01 S/m, σ*_CSF_* 1.654). In all models, we injected electrical current with 1 mA magnitude in LR and AP directions to calculate ***J*** distributions in the brain using FEM for numerically solving the Laplace equation for the electric potential (Saturnino et al., 2019a). The current-induced ***B*** distributions were determined from the current density by applying the Biot–Savart law using the Fast Fourier Transform (Yazdanian et al., 2020). In particular, the z-component of the current-induced magnetic flux density *B_z_* is given by

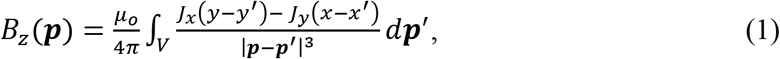

where ***p*** = [*x,y,z*] is the position at which *B_z_* is evaluated, *V* denotes the head model and ***p’*** = [*x′, y′, z′*] are positions inside *V*. ***J*** = [*J_x_,J_y_,J_z_*] is the current density in the head model and constant *μ_o_* is the magnetic permeability of free space.

**Figure 1.**
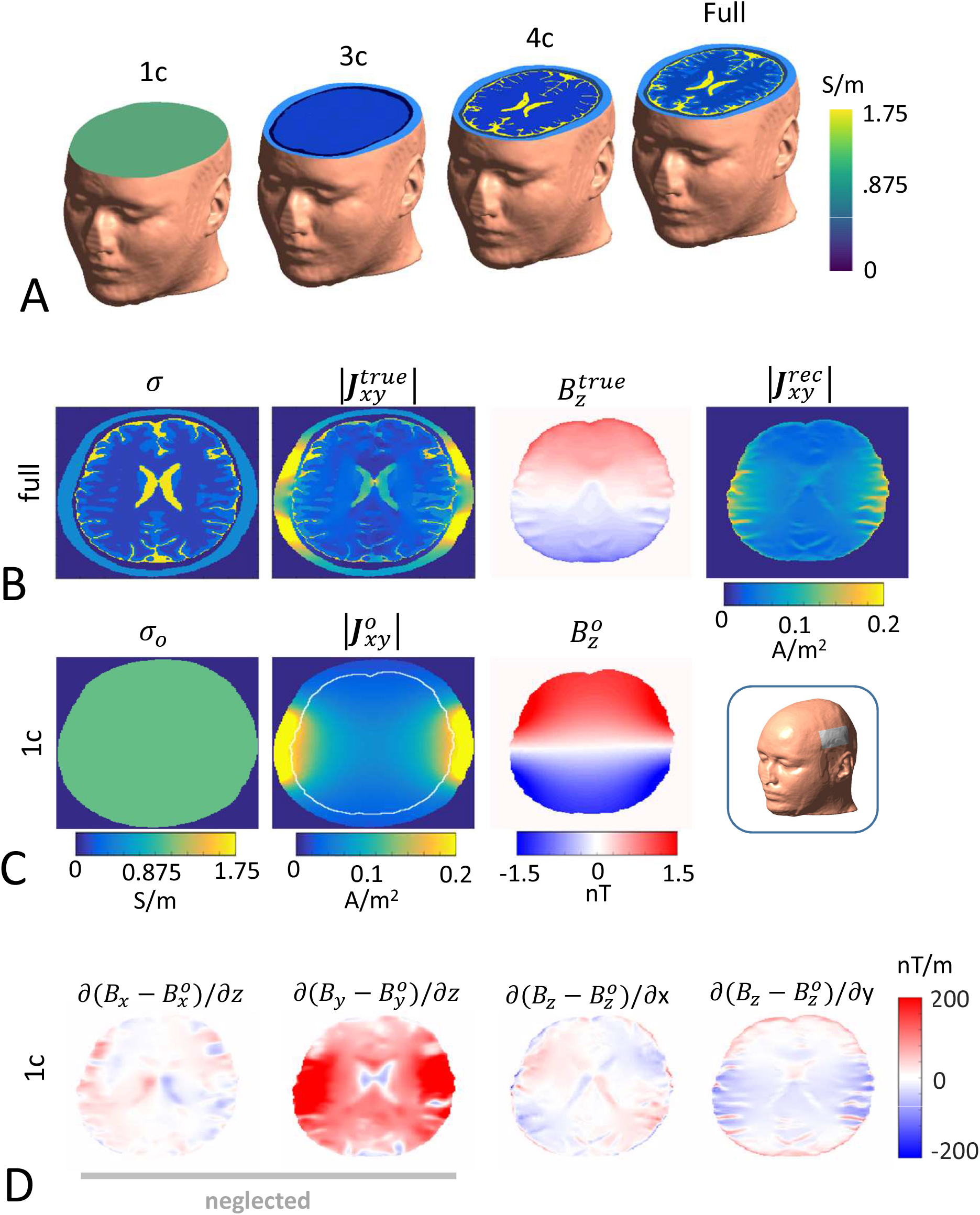
Current density reconstruction for a LR current injection. (A) Volume conductor models of the head with different levels of anatomical detail (1c, 3c, 4c, and full models). The results are shown for an axial slice of these models. (B) The conductivity distribution σ of the full model used as ground truth and the corresponding simulated 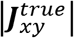 and 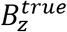 distributions are shown in columns 1-3. The current density distribution 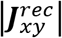 that was reconstructed from 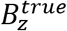 by the projected current density algorithm is shown in column 4. (C) Conductivity, current density and magnetic field distributions for the homogenous (1c) head model that was used in the projected current density algorithm. (D) Visualization of the terms that are neglected and included, respectively, in the projected current density algorithm (see eq. 3) for the 1c head model.

In addition, we aimed to test how much the projected current density ***J**^rec^*, reconstructed in the imaging slice using the projected current density algorithm (equation (2) below), changes the current-induced *B_z_* distributions. For that, we substituted the calculated ***J*** distributions by ***J**^rec^* at the positions corresponding to CSF and brain in the imaging slice before determining the current-induced ***B*** distribution.

### Projected Current Density Algorithm

The projected current density algorithm introduced by Jeong et al (2014) reconstructs a current density distribution ***J**^rec^* from a *B_z_* image

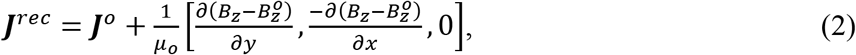

using simulated current densities 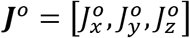 and magnetic flux densities 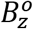 that are based on a volume conductor model of the subject. In the original form of the algorithm, ***J**^o^* and 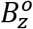 are obtained for a model with a homogeneous conductivity σ*_o_* (1c) and the outer shape of the imaged object (Fig. 1B). The reconstructed current density distribution ***J**^rec^* is then composed of a smooth and curl-free component ***J**^o^* that is superimposed onto a high contrast component based on directional derivatives of 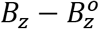. The latter scales with the local difference between measured and simulated magnetic flux densities.

The algorithm can be easily derived from Ampère’s law. The unknown true current density ***J*** that gives rise to the magnetic field ***B*** can be written as:

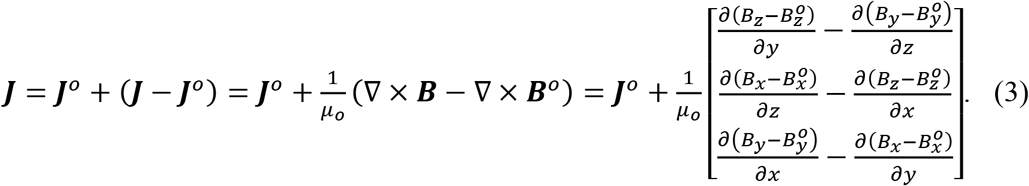

As we only measure *B_z_*, our best possible assumption about the spatial derivatives of the two unknown components of ***B*** are 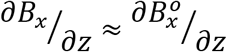 and 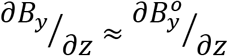. In this case, equation (3) simplifies to equation (2). For example, this assumption will approximately hold in phantoms that is uniform along the z-direction and in which the z-component of the injected current can be kept small.

The algorithm is straightforward to implement and can be applied to single cross-sectional simulated *B_z_* images in case of limited volume coverage. In contrast to the original projected current density method that employs a second-order Laplacian operator (Park et al., 2007), the modified variant proposed by Jeong et al (2014) relies only on first-order spatial derivatives of the *B_z_* image data. This reduces the amplification of high-frequency noise by the derivative operations and potentially improves the quality of the reconstructed current density images from *B_z_* data with lower signal-to-noise ratios (SNR). In addition, the method introduced by Jeong et al (2014) only requires local derivative operations, making it applicable to data from objects that contain large regions with low or no signal such as the human head with the skull and air cavities. On the downside, an error analysis by Jeong et al (2014) suggests that the simplified method will likely suffer from higher reconstruction errors for high SNR *B_z_* images compared to the original approach suggested in (Park et al., 2007).

### Error Metrics

We evaluated the difference between a current density ***J**_xy_* and a reference current density 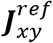 using the relative root mean square (RMS) differences:

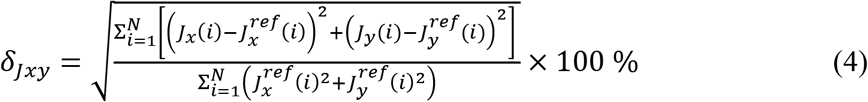

Here, *i* is the pixel index and *N* is the number of pixels inside the measured region composed of WM, GM and CSF. Please note that we restricted the comparison to the x- and y-components of the current flow, as only those influence the z-component of the magnetic flux density *B_z_* measured by MRI and as only the x- and y-components of the current density are changed by the projected current density algorithm. For completeness, we also evaluated the difference between *J_z_* and the z-component of a reference current density 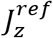 in selected cases as

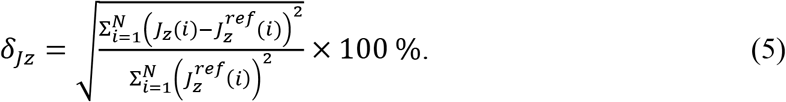

In addition, we quantified the differences between the z-component of the current induced magnetic flux density *B_z_* and a reference case 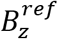 as

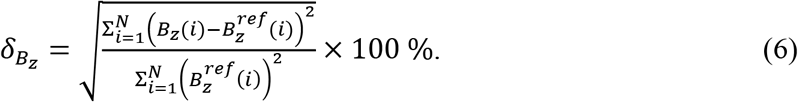

Finally, we calculated relative noise floors (*nf*, the inverse SNR) for the experimentally measured current-induced *B_z_* images as

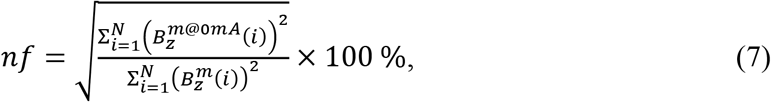

where 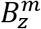 and 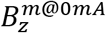 are the measured *B_z_* fields with and without current injection, respectively.

### Optimization of conductivities

As alternative approach to estimate current densities from magnetic flux densities, we optimized the tissue-specific conductivities using the sequential least squares programming (SLSQP) algorithm (Kraft, 1988) with the goal to minimize the error *δ_B_z__* between the simulated magnetic flux density *B_z_* given by the head models and the reference 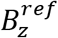. Initially, we used the simulated magnetic flux density of the full model with all tissue types as reference to confirm the stability of the optimization approach. Subsequently, measured *B_z_* images from the human imaging data served as reference. We implemented constraints on the conducitivities to reduce the likelihood of overfitting that could occur, e.g. in case of systematic differences of the volume conductor models to the true head anatomy. For the 3c model, we constrained σ*_scalp_*, σ*_skull_* and σ*_inside_* to the ranges [0.2 < σ*_scalp_*, 0.003 < σ*_skull_* < 0.03, 0.2 < σ*_inside_* < 0.95] S/m. For the 4c model, we used the ranges [0.2 < σ*_scalp_* < 1, 0.003 < σ*_skull_* < 0.03, 1.2 < σ*_CSF_* < 1.9, 0.05 < σ*_brain_* < 0.95] S/m. We also fitted the full model with five conductivities (5c) to human imaging data by using the ranges [0.2 < σ*_scalp_* < 1, 0.003 < σ*_skull_* < 0.03, 1.2 < σ*_CSF_* < 1.9, 0.05 < σ*_WM_* < 0.95, 0.05 < σ*_GM_* < 0.95] S/m. For the 5c model, we imposed σ*_GM_* > σ*_WM_* as an additional constraint. For the 5c model, the ranges were chosen to well include the conductivities reported in studies that measured relatively fresh or live tissue, preferably human at low frequencies (0–100 kHz) near body temperature (Saturnino et al., 2019b, Table 1). The ranges of the combined tissue regions of the 3c and 4c models were chosen correspondingly using volume-weighted averages of the ranges for the tissues of the 5c model. We initialized the optimization algorithm with [*σ_scalp_* = 0.465, *σ_skull_* = 0.01, *σ_inside_* = 0.34] S/m for the 3c model and [*σ_scalp_* = 0.465, *σ_skull_* = 0.01, *σ_brain_* = 0.18] S/m for the 4c model and [*σ_scalp_* = 0.465, σ*_skull_* = 0.01, σ*_CSF_* = 1.654, σ*_WM_* = 0.126, σ*_GM_* = 0.275] S/m for the 5c model.

It is worth noting that without additional knowledge of the total resistance between the two stimulation electrodes, the tissue conductivities are only determined up to a common scaling factor. That is, for a head model with N tissue compartments, only N-1 conductivity *ratios* are uniquely determined via the optimization algorithm. We could have directly optimized N-1 conductivity ratios instead. However, this would have required the choice of an arbitrary “reference” tissue and would have made the use of biologically plausible conductivity ranges as constraints in the optimization difficult.

Applying the optimization algorithm requires the repeated evaluation of *B_z_* for varying conductivities of the head models. This is computationally expensive when it is based on FEM simulations for obtaining the current densitiy ***J*** and subsequent solving of the Biot–Savart law using the Fast-Fourier Transform. Therefore, we employed a non-intrusive generalized polynomial chaos (gPC) expansion for the fast evaluation of *B_z_* as a function of the conductivities (scalp, skull, CSF, GM, WM) (Codecasa et al., 2015; Ghanem et al., 2017; Saturnino et al., 2019b; Wiener, 1938). For the gPC expansions, we used a Jacobian polynomial basis set with a maximum order of 10 and sampled the conductivities by using uniform probability density functions. We selected the conductivity ranges of the gPC expansions slightly larger than the corresponding bounds of the optimization algorithm in order to ensure sufficient accuracy of the *B_z_* fields across the tested conductivity range. We used K-fold cross validation with *K* = 10 for the evaluation of the relative error between the *B_z_* fields obtained from the forward simulations and gPC expansions, and set the relative error of the gPC method to 1%. Training of the gPC converged after ~250 FEM simulations. Using the gPC model, we were then able to reevaluate the *B_z_* fields in the complete head model as a function of the conductivity parameters with a computation time in the order of a few seconds. For further details on our gPC implementation, please refer to (Saturnino et al., 2019b).

### Statistical Testing

Taking the measured current-induced *B_z_* images from the human imaging data as reference, we assessed the differences *δ_B_z__* (eq. 6) between the simulated and measured *B_z_* data as an indication of the quality of the simulations. In particular, we compared two simulations by calculating 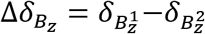, where 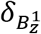 and 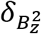 denote the root mean square differences between simulations and measured *B_z_* data. We generated *H_o_* distributions by randomly assigning the voxels of the two simulated *B_z_* images to two “test images” and calculating Δ*δ_B_z__* for these test images. This was repeated 10000 times. We then calculated a one-sided p-value, testing if a more complex model results in smaller differences when compared to the reference data. This was done by assessing the proportion of sampled permutations where the difference was greater than or equal to the observed difference Δ*δ_B_z__*. Significance was then assessed at p=0.01, Bonferroni corrected for the number of comparisons.

In order to systematically compare the 5c, 4c and 3c head models with optimized conductivities, we randomly selected 80% of the imaged slice as input for the optimization algorithm. Then, we calculated the errors 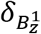 and 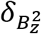 for the remaining 20% of the slice for each of the two head models, and determined the difference 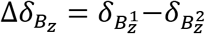, between the simpler 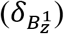 and the more complicated model 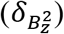. This was repeated 10000 times. Finally, as above, we calculated a onesided p-value by assessing the proportion where the absolute difference was greater than the observed difference Δ*δ_Bz_*, and tested for significance at p=0.01, Bonferroni corrected for the number of comparisons. We used the same procedure to compare the conductivity-optimized 3c head model with the 1c head model, however keeping the conductivity of the 1c model fixed and only repeatedly optimizing the 3c model.

### Data and Code Availability

The original MR data cannot be made publicly available due to privacy restrictions. The largest part of the methods is publicly available via our project homepage www.simnibs.org. This includes the methods for building the head volume conductors from structural MR images, forward simulations of the current densities and current-induced magnetic fields and non-intrusive generalized polynomial chaos expansions for fast re-evaluation of the current-induced magnetic fields in case of changing conductivities. The scripts for statistical testing were developed to address our immediate research needs, but not as easy-to-use software for dissemination and will therefore not be made publicly available at the current time point.

## RESULTS

In the following, we start by characterizing the performance of the “standard” projected current density algorithm, using simulated ground truth data as reference. We then assess whether the reconstruction accuracy of the algorithm can be improved by using more realistic volume conductor models, and test how sensitive the reconstructed current density is to errors of the head models when the latter become more detailed. We continue by evaluating the performance of an alternative approach to estimate the current density that is based on optimizing the conductivities of detailed volume conductor models to maximize the fit of the simulated and the reference *B_z_* data. Complementing our initial assessments based on simulated data, we test the two approaches for *B_z_* data obtained from MR imaging of the human brain. In particular, we introduce a hierarchical statistical testing approach as a principled way to test and compare the quality of reconstructed current density images that accounts for the limited signal-to-noise ratio of measured *B_z_* data and the fact that the ground truth of the current density is unknown in reality.

### Projected Current Density Algorithm based on a Homogenous (1c) Head Model

Figure 1A shows the conductivity distribution *σ* in an axial slice through a realistic head model (first column) and the simulated current density distribution 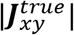 for a left-to-right (LR) current injection (second column). The z-component 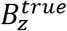 of the magnetic field arising from the injected current flow is shown in the third column. In practice, MRI measures *B_z_* only inside CSF and the brain, which is considered here by applying a corresponding mask. Applying the standard projected current density algorithm to the *B_z_* distribution results in the reconstructed current distribution 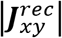 shown in the fourth column of Figure 1A. It is obvious that the reconstructed current density differs strongly from the true simulated current density distribution inside the cranial cavity both in terms of its spatial pattern and average strength (R^2^=0.22 between 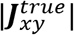 and 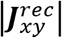; average strengths: 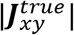 0.039 A/m^2^, 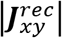 0.058 A/m^2^). In particular, 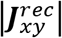 fails to capture the spatial details of the current flow, even those that originate from large anatomical structures such as the ventricles. In contrast, the spatial pattern of 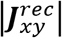 is more similar to the current flow 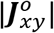 inside a homogeneous conductor (Figure 1B, second column; R^2^=0.68 between 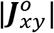 and 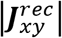, average strength of 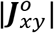: 0.068 A/m^2^). This shows that the results of the projected current density algorithm are dominated by the simulated current density ***J**^o^* (eq. 2), and are less influenced by the actually measured *B_z_* data. The reason for this low performance gets apparent when visualizing the terms that are neglected in the projected current density algorithm (see eq. 3) as they involve the non-measurable *B_x_* and *B_y_* components of the magnetic field (first and second column of Fig 1C). For the simulated head anatomy, the neglected terms are as strong and even stronger than the included terms that depend on the measured *B_z_* component (third and fourth column of Fig 1C), which prevents a reasonable reconstruction of the true current density distribution.

The projected current density algorithm shows similar performance for an anterior-to-posterior (AP) current direction (rows one and two of Fig. S1A). Very similar results were also obtained when using a head model as ground truth that incorporated anisotropic brain conductivity, derived from diffusion MR data (Fig. S2). These control results confirm the limited applicability of the algorithm for realistic head anatomies. In contrast, the algorithm performs better for a volume conductor that is uniform along the z-axis, as the neglected terms are close to zero in this case (row three in Fig. S3). For this simplified “head” model, 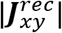 captures the spatial details of 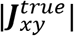, especially throughout the ventricles (R^2^=0.63 between 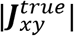 and 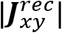 average strengths: 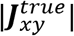 0.035 A/m^2^, 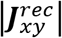 0.032 A/m^2^). The remaining differences show mostly up as blurring, caused by a limited numerical accuracy of the gradient calculations in equation (2). This effect has also been observed in previous studies using simplified phantoms (Sajib et al., 2012).

### Projected Current Density Algorithm based on Inhomogeneous Head Models

Using more realistic volume conductor models of the head can help to make the simulated ***J**^o^* more similar to the true current density. By that, the strength of the terms that are neglected in the projected current density algorithm will also be reduced. In the following, we tested whether this helps to improve the performance of the algorithm. For that, the reconstructed current densities were compared when using three different head models to calculate ***J**^o^*: 1) A model with a single compartment (1c) as used in the original algorithm. 2) A model with three compartments corresponding to scalp, skull and the cranial cavity (3c in Fig. 2B). 3) A model with four compartments corresponding to scalp, skull, CSF and the brain (4c in Fig. 2B). Please see the Methods section for the choice of the conductivities for the 3c and 4c models.

**Figure 2.**
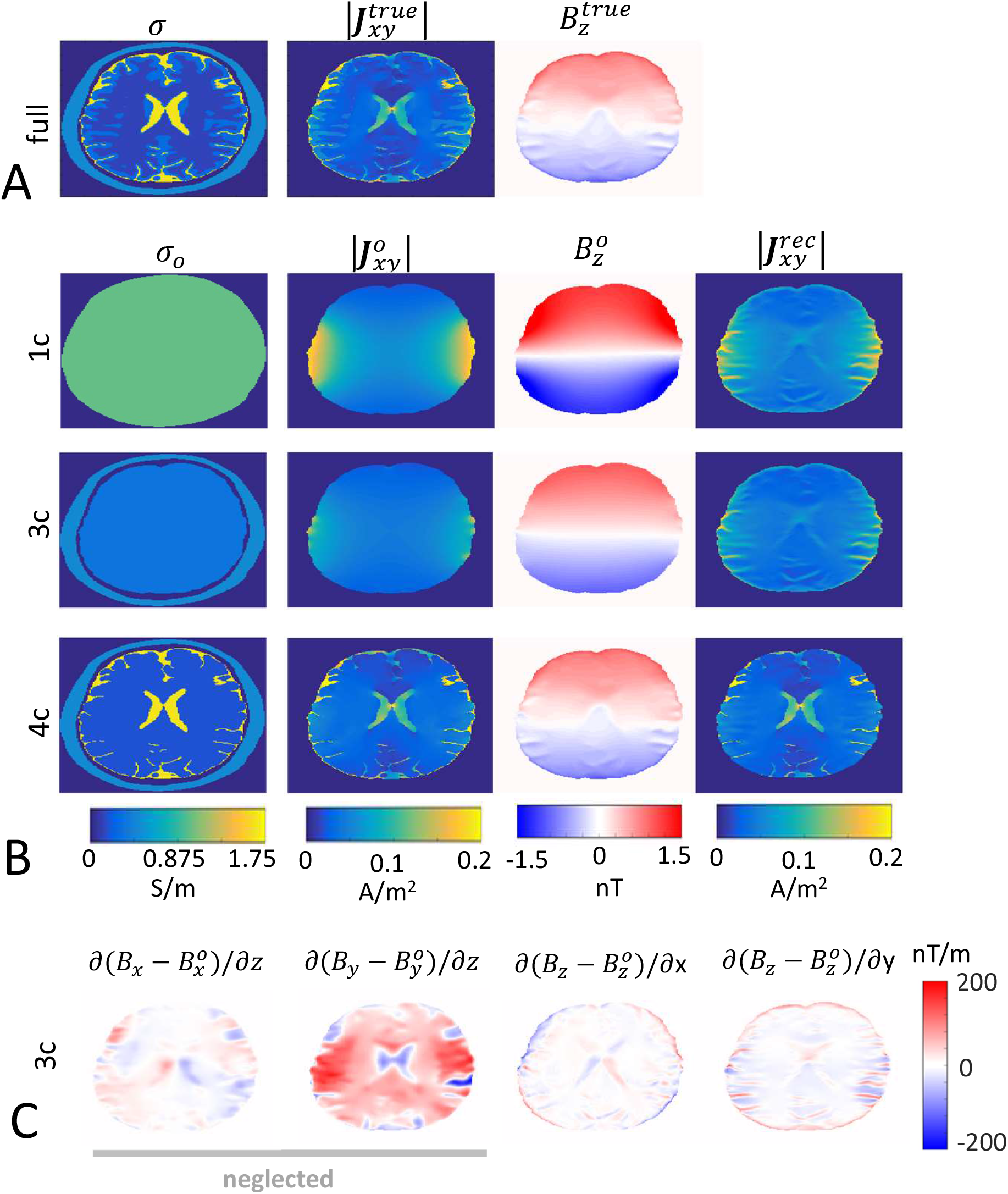
Current density reconstructions based on different head models for a LR current injection. (A) The conductivity distribution *σ* of the full model used as ground truth and the corresponding simulated 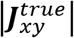 and 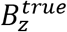 distributions. (B) Results for the 1c, 3c and 4c head models. Columns 1-3 show the conductivity distributions and the simulated 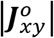 and 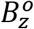 distributions. Column 4 depicts the current densities 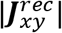 that were reconstructed by the projected current density algorithm with the corresponding head model. (C) Visualization of the terms that are neglected and included, respectively, in the projected current density algorithm based on the 3c head model.

Expectedly, the reconstructed current density (4^th^ column in Fig. 2B) approaches the true current density distribution (Fig. 2A) when the head models get closer to the full model used as reference. For example, the current density in the brain is clearly overestimated close to the electrodes when using the 1c model that does not account for the “shielding” effects of the low-conductive skull. This effect is mostly corrected by using the 3c model, but only the 4c model also achieves a reasonable estimation of the current density around the ventricles and in the sulci. Evaluation of the RMS errors *δ_Jxy_* (eq. 4) for the 1c, 3c and 4c models (Fig. 3A) reveals that most of the improvement resulted from the better models used to calculate ***J**^o^*, while the projected current density algorithm causes only little additional improvement. The underlying reason is that the neglected terms are still very strong even for the more detailed models (Fig. 2C shows all terms for the 3c model as example). For the 3c model, the algorithm reconstructs some of the details of the current flow in the brain that are missing in the underlying 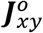 distribution (second row of Fig. 2B). However, this results only in a small overall improvement in *δ_Jxy_* (Fig. 3A). A visual inspection of the spatial distribution of the reconstruction error revealed that the projected current density algorithm decreased the error in many parts of the brain, but also resulted in localized increases at several positions, in particular around the sulci (data not shown). For confirmation, an anterior-to-posterior current direction was also tested (Figs. S1 and S4), giving similar results.

**Figure 3.**
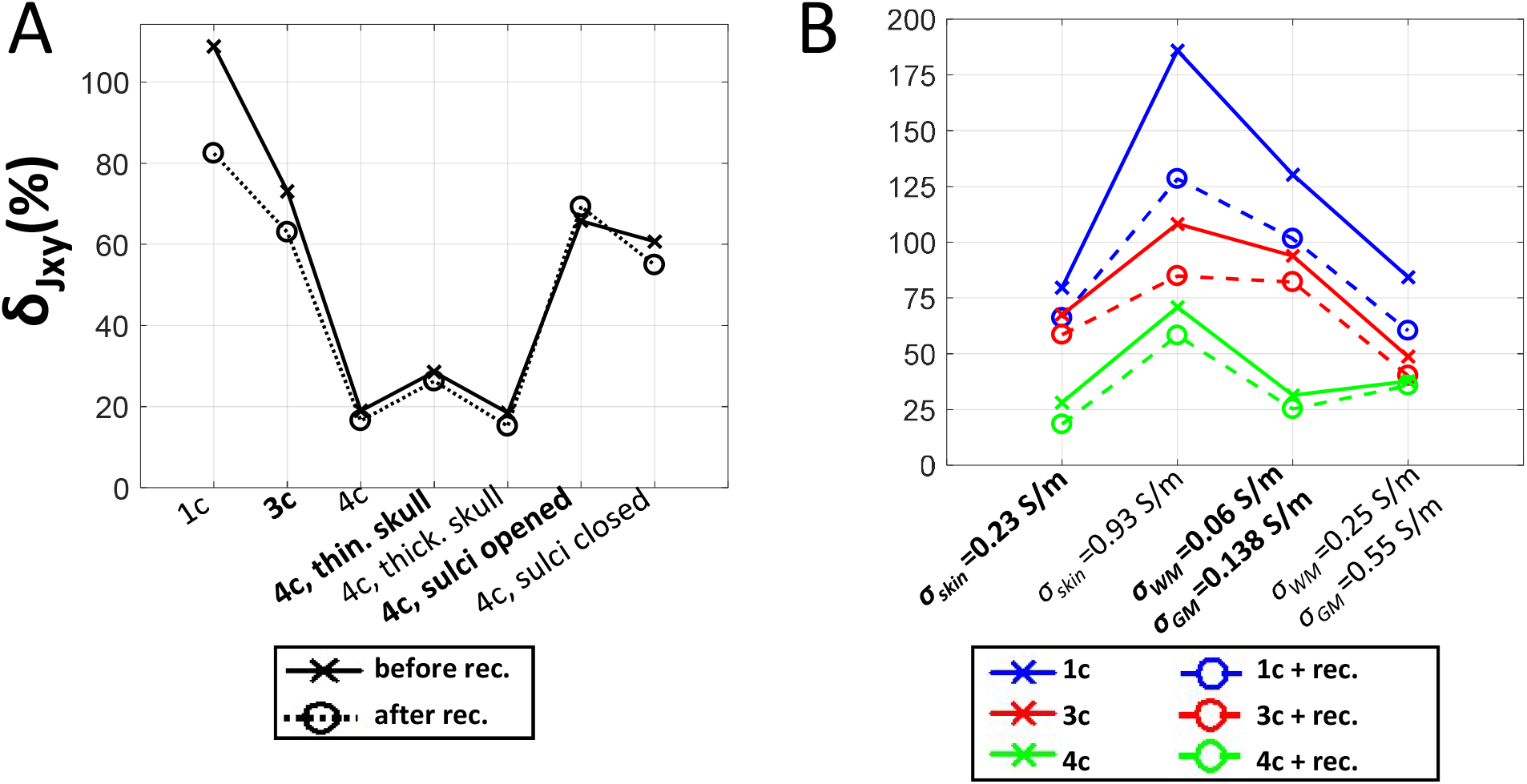
RMS errors *δ_Jxy_* of different head models for a LR current injection, with and without applying the projected current density algorithm. Solid and dashed lines represent the *δ_Jxy_* values for the 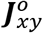 and 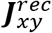 distributions, respectively. (A) Dependence of *δ_Jxy_* on the anatomical level of detail of the head model (1c, 3c and 4c models) and on segmentation errors of the 4c model. (B) Dependence of *δ_Jxy_* on variations of the conductivity of the full model that was used as ground truth.

While using more realistic head models seems straightforward to reduce the reconstruction error, it is important to note that these models still include errors, e.g. due to inaccurate segmentations of the structural MR images or incorrectly chosen ohmic tissue conductivities which deviate from their unknown ground truth. That is, more detailed head models will result in current density distributions appearing realistic with more spatial high frequency content, but which are not necessarily closer to the real current density distribution. In the following, it is therefore tested how robustly improvements can be obtained when the head model for the ***J**^o^* simulations differs from the real volume conductor and whether the projected current density algorithm is helpful to correct errors introduced by deviations of the head models from the ground truth.

Fig. 3A shows the RMS errors *δ_Jxy_* for selected variations of the anatomy of the 4c model, namely overall thinner and thicker skulls (affecting the amount of current entering the cranial cavity) and opened and closed sulci (affecting the amount of well-conductive CSF in the sulci and by that the local current flow distribution). While varying the skull thickness changed the current flow only little, modifying the sulci increased the error almost to the level of the 3c model (Fig. S5 shows the corresponding 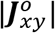 and 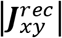). Fig. 3B depicts the RMS errors for the homogenous, 3c and 4c models when varying selected conductivities of the full model that was used as ground truth. The scalp conductivity of the full model was set to half and double of the standard value, and the conductivities of GM and WM were halved and doubled. In all cases, the 3c and 4c models maintained a better performance than the 1c model. Both for the tested variations of anatomy and conductivity, the projected current density algorithm achieved no or only moderate improvements. In particular, changing to anatomically more accurate head models resulted in stronger improvements than those that could be obtained by using the algorithm.

### Optimization of the Tissue Conductivities of the Inhomogeneous Head Models

The above results indicate that the fit between the estimated and true current densities can be improved more by choosing an appropriate volume conductor model than by applying the projected current density algorithm. A straightforward approach to estimate the current density distribution is thus to use the measured *B_z_* data to optimize the tissue conductivities of anatomically detailed volume conductor models that are determined via the segmentation of structural MR data. The following section provides a basic proof-of-concept that this optimization approach can successfully account for conductivity changes of the full model used as ground truth. The performance for volume conductor models that differ in the amount of anatomical detail is also tested and it is assessed how the presence of inaccurate segmentations impacts reconstruction performance.

It can be exemplarily seen in Fig. 2A and B that the similarity of the estimated and true current density distributions 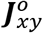 and 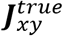 co-vary with the similarity between 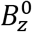 and 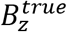 when changing between the 1c, 3c and 4c models. This is also revealed in the corresponding co-variation between the RMS errors *δ_Jxy_* and *δ_B_z__* in Fig. 4A and C (solid lines – 1c, 3c and 4c). In addition, when varying the conductivities of the full model that serves as ground truth (solid lines in Figure 4B and D), both errors show similar dependences on the conductivity and the model (R^2^=0.86 between *δ_Jxy_* and *δ_B_z__* for the non-optimized cases). That is, the errors in ***J**_xy_* relative to its ground truth are well reflected in the errors in *B_z_*. This favorable behavior of *B_z_* occurs as, by the Biot-Savart law, *B_z_* is fully determined by the three-dimensional distribution of ***J**_xy_*, with the impact of the current density scaling with distance to the measured *B_z_* (eq. 1). It is interesting to note that, by Ampère’s law, ***J**_xy_* is determined by the spatial changes of all components of the magnetic field rather than only *B_z_* (eq. 3). This explains why the projected current density algorithm can have low performance despite a close relation between ***J**_xy_* and *B_z_*.

**Figure 4.**
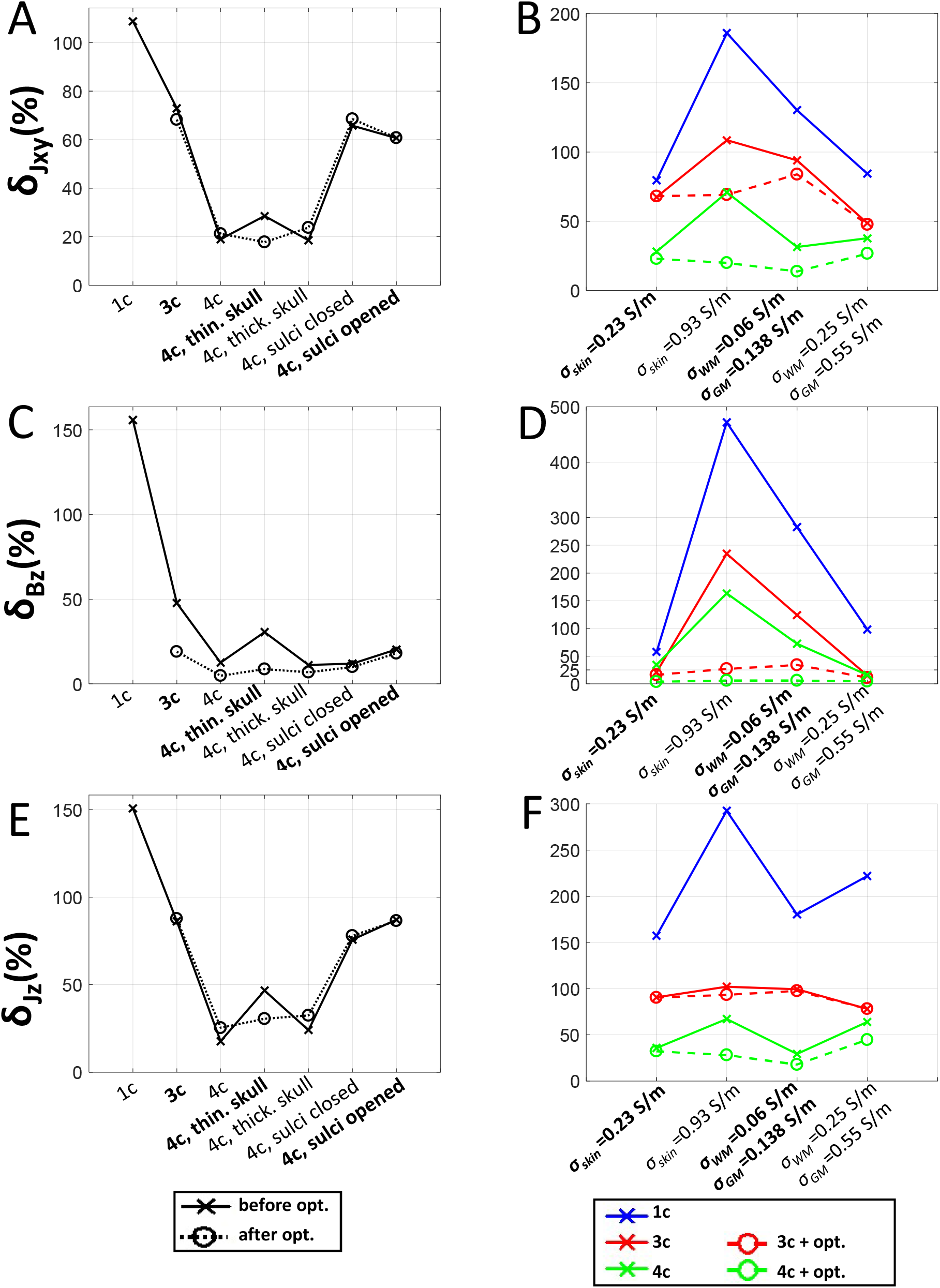
RMS errors *δ_Jxy_, δ_Bz_* and *δ_Jz_* of different head models for a LR current injection, with and without optimized conductivities. Solid and dashed lines correspond to the error values obtained for the models with literature and optimized conductivities, respectively. The first column (A, C & E) shows the dependence of *δ_Jxy_, δ_Bz_* and *δ_Jz_* on the anatomical level of detail of the head model (1c, 3c and 4c models) and on segmentation errors of the 4c model. The second column (B, D & F) shows the dependence of *δ_Jxy_, δ_Bz_* and *δ_Jz_* on variations of the conductivity of the full model that was used as ground truth.

The demonstrated relation between *δ_Jxy_* and *δ_B_z__* suggests that minimizing the RMS errors for *B_z_* by optimizing the conductivities of the head models might be a useful approach to improve the similarity between the simulated ***J**_xy_* and the ground truth. Indeed, optimizing the tissue conductivities of the 3c and 4c models to minimize *δ_B_z__* consistently improves the fit for ***J**_xy_* (dashed lines in Fig. 4B and D, R^2^=0.86 between *δ_Jxy_* and *δ_B_z__* for the optimized cases). Expectedly, the achievable improvement scales with the level of detail of the head model. Rather trivially, the errors approach zero when the full head model is used for the optimizations (data not shown). In general, optimization reduces *δ_B_z__* more than *δ_Jxy_* when compared to the RMS errors of the nonoptimized cases. Still, also after optimization, the remaining *δ_B_z__* reflect well the relative differences between the *δ_Jxy_* of the different head models, making the former a useful parameter for model comparison.

Optimizing the tissue conductivities in the presence of anatomical inaccuracies hardly changes *δ_Jxy_* (dashed line in Fig. 4A) although it reduces *δ_B_z__* for all cases (R^2^=0.61 between *δ_Jxy_* and *δ_B_z__* for the optimized cases). This indicates that constraining the optimization to a few conductivities which affect the current flow globally reduces the risk of overfitting. However, it also shows that accurate segmentation of the brain anatomy from the MR images is required to ensure accurate estimations of local details of the current density distribution via this approach. In particular, also for the non-optimized cases, changing the sulcal structure of the 4c model affects ***J**_xy_*, but hardly changes *B_z_* (solid lines in Figure 4A and C). This indicates that *B_z_* is not a sensitive marker of localized errors in the estimated current density distribution.

For confirmation, an anterior-to-posterior current direction was also tested (Fig. S6), giving similar results. The error for *J_z_* is shown in Fig. 4E and F for completeness. Even though the relationship between *J_z_* and *B_z_* is only indirect, it follows the same general dependence on the head model and conductivities as observed for ***J**_xy_* and *B_z_*. As an alternative to optimizing the model conductivities, Kwon and colleagues (2016) introduced an iterative version of the projected current density algorithm that, combined with an anatomically detailed head model, achieved good reconstruction accuracy for simulated ground truth data. However, our initial tests suggest that stable convergence depends on the availability of *B_z_* data for the complete head volume including scalp and skull (Supplementary Material B).

### Hierarchical Model Selection based on Statistical Testing

The results so far indicate that the tested projected current density algorithm has limited capabilities to reconstruct the current density distribution for the complex anatomy of the human head. Instead, a better estimation of the true current density distribution is achievable by using detailed volume conductor models that are reconstructed from structural MR scans and optimizing their conductivities based on the measured *B_z_* data. In contrast to the above tests that used simulated *B_z_* data also as ground truth, measured *B_z_* data contains noise. Therefore, the difference between the simulated and measured *B_z_* fields would not fully disappear even if the head model was perfect. In order to quantify the quality of current density distributions calculated via head models in practice, the difference between the simulated and measured *B_z_* field thus has to be compared to the expected noise level of the *B_z_* measurements, for example obtained in independent control measurements without current injection. Specifically, when optimizing model parameters such as the conductivity values, this comparison can help to identify cases of overfitting in which the remaining difference can be lower than the expected noise level. This is particularly relevant for models with many free parameters. Finally, as the ground truth with regards to the conductivity distribution is unknown, it is important to demonstrate that the current density distribution calculated via a detailed volume conductor model results in *B_z_* data that is closer to the measured *B_z_* data when compared to the current density distribution obtained for an anatomically simpler model. This ensures that increasing the complexity of the head model indeed helps to improve the approximation of the unknown true current density distribution.

In the following, rigorous statistical testing is employed to assess the quality of current density estimates for measured *B_z_* data for two current injection directions in two participants. Initially focusing on the current density estimates obtained for head models with pre-defined conductivities taken from literature, permutation testing is used to assess the difference between the models (see the Methods section for details). Figure 5A shows the measured *B_z_* data of two electrode montages (LR and AP) for one participant, and Figure 5B and 5C depict the simulated data and the remaining difference. The results also confirm that the current density estimated via the 4c model explains the measured *B_z_* significantly better than that of simpler models (Fig. 5D) (Δ = 77.0 % and Δ = 10.7 % between 4c and 3c models for LR and AP, *p* ≤ 0.01 corrected). This also holds for the current density estimated via the standard projected current density algorithm (“1c+rec.” in Fig. 5), which achieves moderate improvements in *δ_B_z__* compared to the 1c head model (Δ=25.3 % and Δ=8.8 % for LR and AP, *p* ≤ 0.01 corrected). As the projected current density algorithm changes the current density only within the brain and CSF area of the imaged slice, the moderate differences in *δ_Bz_* are partly caused by the need to use ***J**^o^* of the 1c model in the remaining parts of the head volume for calculating *B_z_*. In order to ensure a fair comparison to the 3c model, we use the same approach when determining *B_z_* of the 3c model (indicated by the squares in Fig. 5D), still revealing statistically significant differences of 11.8 % and 3.1 % for the LR and AP injections, respectively. The results also show that the current density estimated using the 5c model explains the measured *B_z_* significantly better than the 4c model for the LR injection (Δ=14.6 %, *p* ≤ 0.01 corrected), whereas we do not observe a statistically significant difference for the AP injection (Fig. 5D). For the AP montage, the impact of the ventricles on the *B_z_* distribution in the image center are visible in both the measured *B_z_* data and the results of the 4c and 5c models, but are absent for the simpler models. However, even for the 4c and 5c models, the remaining differences are above the expected noise floor seen for the control measurement for 0 mA. In addition, they exhibit clear spatial patterns, suggesting that the 4c and 5c models do not fully explain all aspects of the underlying current density distribution.

**Figure 5.**
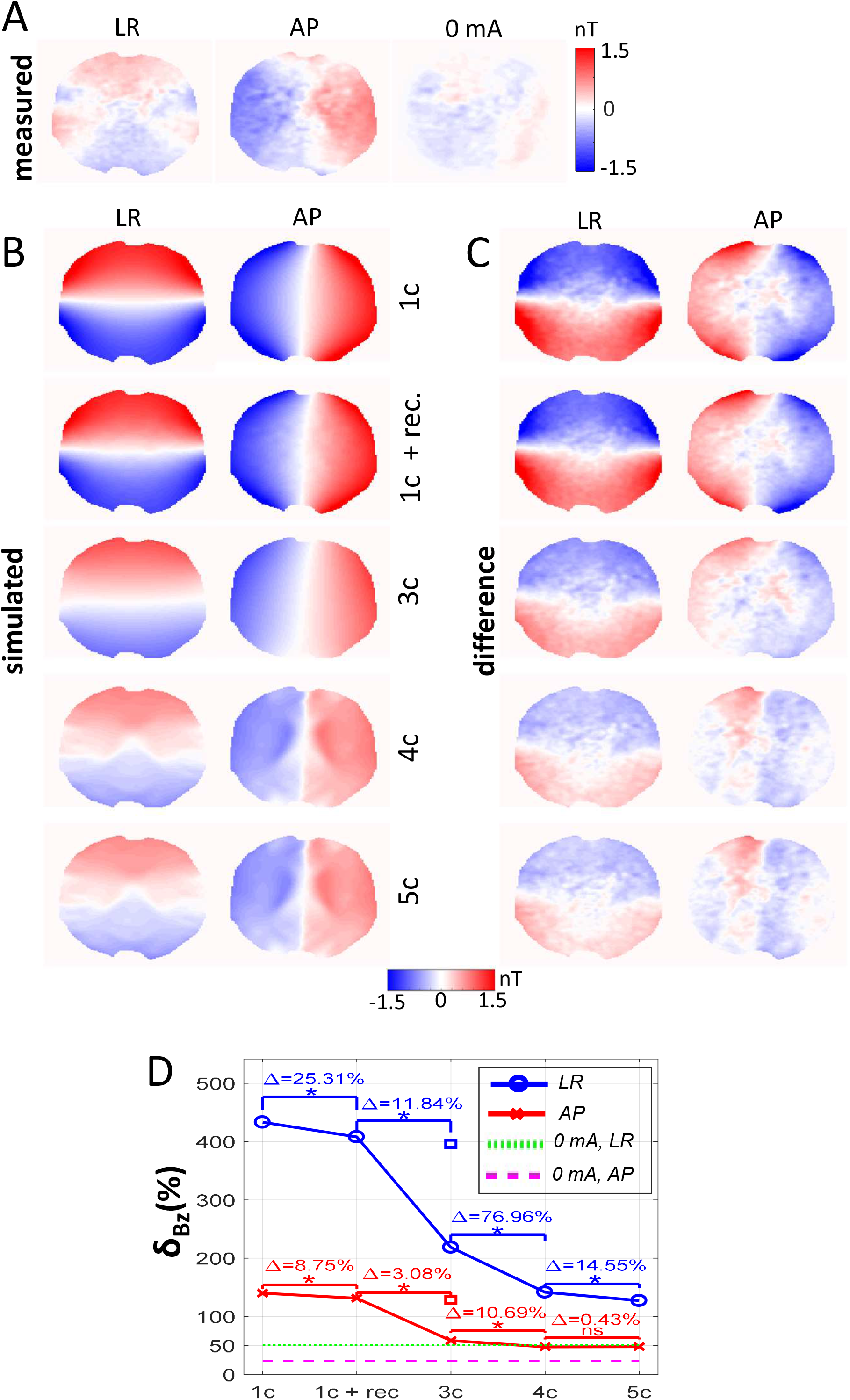
Experimental *B_z_* measurements and the corresponding simulated *B_z_* distributions for the first subject. The conductivity values of the 3c, 4c and 5c models were chosen from literature values, using the procedure described in Methods section. (A) Measured *B_z_* distributions with and without 1 mA current injections in LR and AP directions. (B) Simulated *B_z_* distributions obtained for the 1c, 3c, 4c and 5c models and for the projected current density reconstruction based on the 1c model (1c+rec.). For the latter, *B_z_* was calculated by using the reconstructed current density ***J**^rec^* in the imaged slice and ***J**^o^* of the homogenous head model in the rest of the head volume. (C) Differences between the measured and simulated *B_z_* distributions. (D) Dependence of *δ_Bz_* on the model. Asterisks (*) indicate significant differences between the models at *p* ≤ 0.01, Bonferroni corrected for 8 comparisons. The squares (□) represent *δ_Bz_* values obtained for *B_z_* distributions reconstructed using ***J**^o^* of the 3c model in the imaged slice and ***J**^o^* of the homogenous model in the rest of the head volume. This was done for a fair comparison of the 1c+rec. results with the 3c model, as the reconstructed current density algorithm only changes the current distribution in the imaged slice and the change in *B_z_* is correspondingly lower. The dashed green and purple lines represent the noise floors *nf* (eq. 7). The *δ_Bz_* difference between the 4c and 5c models of the AP case is non-significant (ns).

Fig. 7A shows the corresponding current density estimates for the different models. At the SNR of the measured *B_z_*, the projected current density algorithm (1c model) does not recover details of the inhomogeneous current flow pattern that are expected to occur due to the different conductivities between the brain and CSF. Also, the 3c model does not reconstruct any details of the inhomogeneous current flow pattern, but still explains the measured *B_z_* field better because it corrects for the impact of the low-conductive skull on the average strength of the current flow in the cranial cavity. The 4c and 5c models improve the fit to the measured *B_z_* field by accounting for the spatially varying conductivities in the cranial cavity. The results for the second subject are shown in Figs. S7 and S9A, confirming the above observations.

### Hierarchical Model Selection for Models with Optimized Conductivities

The *B_z_* fields of the 3c, 4c and 5c models with optimized conductivities are shown in Fig. 6B for the first subject, and the remaining differences to the measured fields are shown in Fig. 6C. Compared to the models using literature values for the conductivities, optimization reduced the average strength of the simulated *B_z_* fields. This decrease is a main cause of the largely improved *δ_Bz_* of the 3c, 4c and 5c models compared to the 1c case (Fig. 6D vs 5D), and is correspondingly reflected in the on average lower current densities of the optimized 3c, 4c and 5c models in Fig. 7B compared to their counterparts in Fig. 7A. In addition, the optimized *B_z_* fields in Fig. 6B also better reflect the measured spatial variations of the simulated *B_z_* fields close to the electrodes in particular for the LR injections.

**Figure 6.**
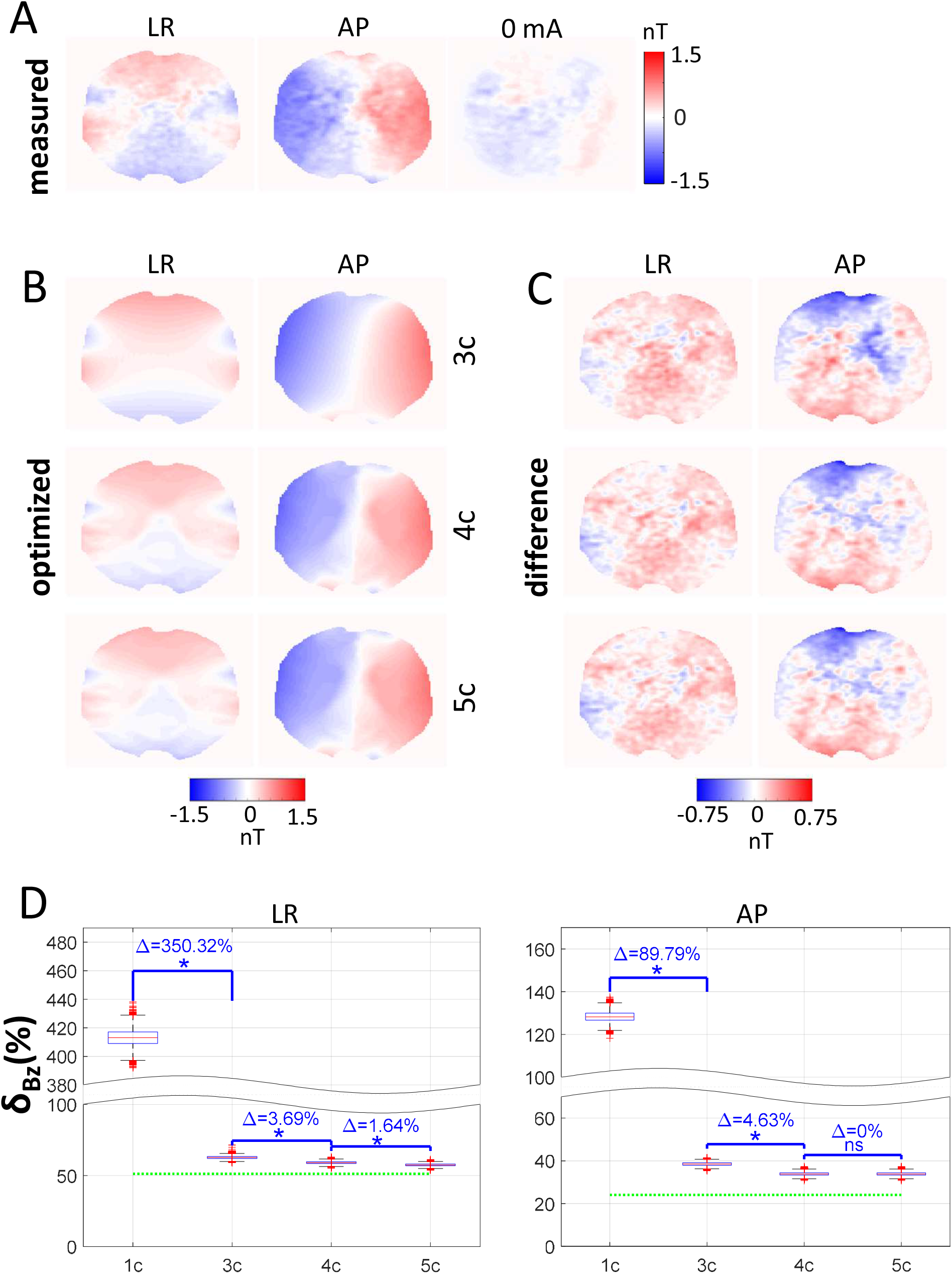
Results for the 3c, 4c and 5c models with optimized conductivities for the first subject, determined by minimizing the difference to the measured *B_z_* distributions. (A) Measured *B_z_* distributions with and without 1 mA current injections. (B) Optimized *B_z_* distributions for the 3c, 4c and 5c models, corresponding to the median *δ_Bz_* shown in D. (C) Difference between the measured and optimized *B_z_* distributions. (D) Dependence of *δ_Bz_* on the model. The red lines indicate the median, the boxes mark the 25th and 75th percentiles and the whisker lengths is set to 1.5 times the interquartile range. Values outside that range are marked as outliers. Blue asterisks (*) indicate significant differences between the models at *p* ≤ 0.01, Bonferroni corrected for 6 comparisons. The dashed green lines indicate the noise floors.

**Figure 7.**
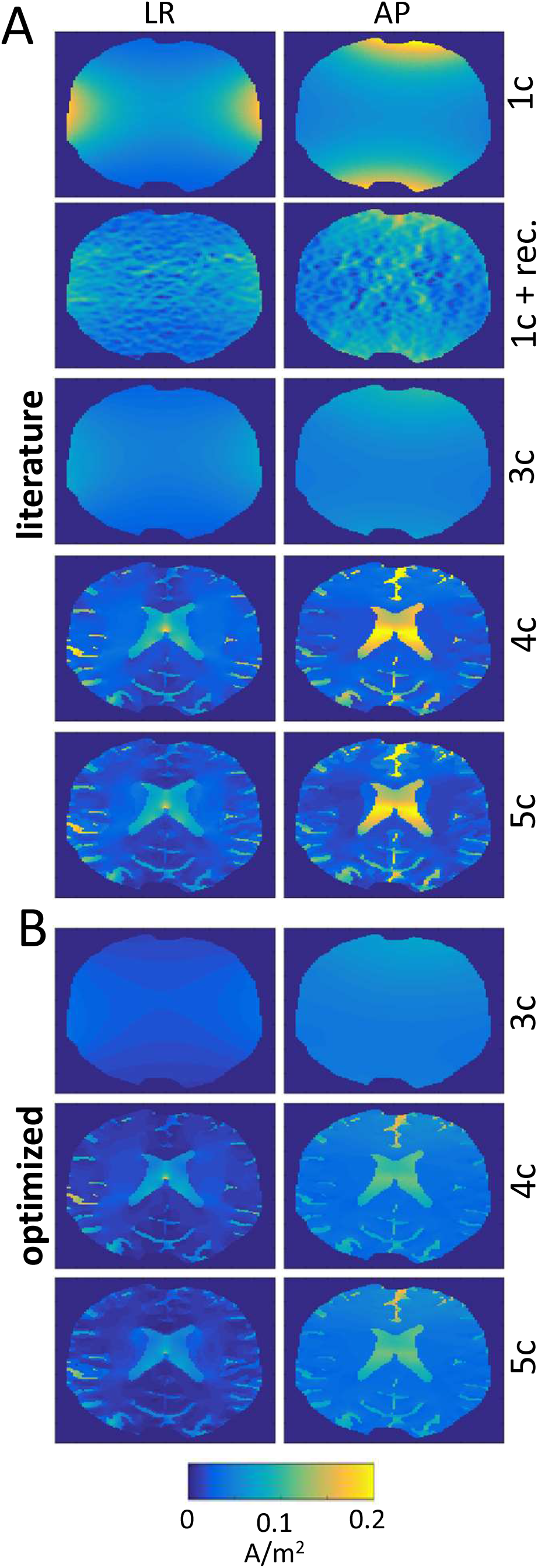
Estimated current density distributions |***J**_xy_*| for the first subject. (A) Results for the 1c, 3c, 4c and 5c models based on literature conductivities. Also, the results for the projected current density reconstruction based on the 1c model are shown (1c+rec.). The corresponding *B_z_* fields are shown in Fig. 5. (B) Results for the 3c, 4c and 5c models based on optimized conductivities. The corresponding *B_z_* fields are shown in Fig. 6.

The improvement of *δ_B_z__* for the 4c versus 3c model is statistically significant for both LR and AP injections. Increasing the level of detail of the model further from 4c to 5c results in statistically significant improvements for the LR injection, but not in case of the AP injection (Fig. 6D). The *B_z_* fields obtained for the 3c, 4c and 5c models result in residuals with similar spatial patterns (Fig. 6C). However, visual inspection of the 4c and 5c models with the 3c model for the AP injection reveals localized variations of the simulated *B_z_* fields in the middle of the brain due to the CSF-filled ventricles. The peak absolute differences in this region are in the order of 0.3 nT, which is similar to the differences of the simulated *B_z_* field of the 3c model and the measured *B_z_* field in that region. These findings indicate the minimal measurement sensitivity that is required to reliably resolve *B_z_* variations caused by large brain structures for a current injection of 1 mA baseline-to-peak. Similar effects of the CSF-filled ventricles are also visible for the AP injection for the second subject in in S8B (S9B shows the corresponding current densities).

While optimization strongly reduces the differences between simulated and measured *B_z_* fields, the remaining differences still show spatial patterns that are above the strength of the noise floor for the 0 mA results. This indicates that also the 4c and 5c models with optimized conductivities do not account for all aspects of the measured *B_z_* field. The optimized conductivities are shown in Fig. S10 for completeness.

## DISCUSSION

Using simulated and measured data, we demonstrated that the tested variant of the projected current density algorithm (Jeong et al., 2014) achieves only coarse reconstructions of the current density distribution in the human brain, while working as expected for artificial geometries that are uniform along the z-direction. The algorithm neglects the components of Ampère’s law (eq. 3) that depend on the *B_x_* and *B_y_* components of the current-induced magnetic field. These components are weak in geometries that are uniform along the z-direction, but can be dominant in case of the human head. This might also explain why the projected current density algorithm achieved little additional improvements when we tested it in combination with more detailed head models instead of a single compartment model. In contrast, increasing the anatomical detail of the head model alone already had a large effect on the accuracy of the estimated current densities. In our tests with simulated ground truth data, this still held when the head model suffered from uncertainties of the ohmic tissue conductivities or had a limited segmentation accuracy. We confirmed that these conclusions derived from simulated ground truth data also apply to measured data: Current densities calculated with a 3-compartment model and tissue conductivities taken from literature explained measured *B_z_* images significantly better than current densities reconstructed using the standard projected current density algorithm.

Our results demonstrate the importance of formal statistical testing for systematic comparisons of volume conductor models that have different levels of anatomical detail. Statistical testing within a hierarchical model selection framework ensured that more detailed models did indeed provide more accurate estimations of the unknown true current density. Expanding the framework beyond models with fixed conductivities, we also demonstrated that the framework allows for the systematic assessment of current density estimates derived from volume conductor models that use optimized tissue conductivities based on the measured *B_z_* images. Complemented by results for simulated ground truth data that showed a stable convergence towards the true current densities with varying tissue conductivities for the employed optimization approach, these findings suggest that using the measured *B_z_* images to fit the tissue conductivities of personalized volume conductor models might be a valuable approach in order to estimate the unknown true current density.

It is worth noting that the possibility to distinguish between the quality of different volume conductor models using measured *B_z_* data and statistical testing also scales with the measurement SNR. Specifically, differences between models will occur as non-significant when the corresponding changes of the *B_z_* data are below the noise levels. While a systematic evaluation of the required SNR levels is outside the scope of this study, our results indicate that a sensitivity of around 0.3 nT is required for a current injection of 1 mA in order to reveal the conductivity contrast caused between the ventricles and surrounding brain areas.

### Relation to Prior Studies

Our findings show that the blurriness and lack of detail of current density reconstructions for human in-vivo data do not merely result from a low SNR of the *B_z_* images (Chauhan et al., 2018; Göksu et al., 2018; Kasinadhuni et al., 2017) and a lack of accounting for the magnetic stray fields cause by the electrode cables in two of the studies (Göksu et al., 2019). Rather, they are also due to a limited applicability of the reconstruction algorithm to the human head anatomy. As the reconstructed current density is biased towards the smooth current flow occurring in a volume conductor with homogenous conductivity (Fig. 1A&B), it is likely that the similarity between the current density distributions reconstructed from measured *B_z_* data of different persons will be artificially increased, camouflaging interindividual differences. This effect also explains why in our prior study the fit between current densities reconstructed from simulated and measured *B_z_* data were consistently better than the fit between the simulated and measured *B_z_* data itself (Göksu et al., 2018a). It is likely that also the performance of methods such as DT-MREIT (Jeong et al., 2017), which uses the reconstructed current densities in combination with diffusion tensor images for estimating the conductivities of brain tissues, will be markedly affected by the low accuracy of the spatial patterns and strengths of the reconstructed current densities, leading to biased conductivity estimates. We only assessed a specific variant of the projected current density method (Jeong et al., 2014) so that our findings will not necessarily generalize to other current density or conductivity reconstruction methods. However, they point towards the need to carefully ensure the desired behavior of reconstruction algorithms when applying them to *B_z_* data of the human head. The lacking uniformity along the z-direction and the lack of data from the scalp and skull regions render the reconstruction problem fundamentally more difficult than for typical MREIT phantoms.

Kwon and colleagues (2016) introduced a method that combines an anatomically detailed head model with an iterative updating procedure to improve the accuracy of the reconstructed current densities by minimizing the difference between calculated and measured *B_z_* data. It showed promising performance on a simulated case in which *B_z_* data were available for the complete head volume. However, our initial tests indicate that it requires further development to be applicable to measured *B_z_* data with limited volume coverage and lack of data from the scalp and skull (Supplementary Material B).

Our approach to optimize conductivities shares similarities with the algorithm proposed in (Gao et al., 2006). Both methods optimize the conductivity values of a few tissue types instead of aiming at a voxel-wise conductivity reconstruction, and both methods use polynomial expansions (here: generalized polynomial chaos expansion; Gao et al.: response surface method) for representing the functional dependence between simulated *B_z_* and conductivity in order to avoid the costly reevaluation of the forward model during conductivity optimization. They differ in the level of detail of the head models and the optimized cost functions. While we chose to minimize the relative root mean square differences between measured and simulated *B_z_*, Gao and colleagues choose a more complex cost function that additionally incorporates the correlation coefficient. While beyond the scope of this study, the impact of the choice of the cost function on the stability and accuracy of the optimization procedure for low-SNR *B_z_* data might be a relevant topic to develop the overall approach further. The largest difference concerns our use of statistical testing for model selection in order to ensure the quality of the estimated current density distribution despite the lack of a ground truth in case of measured *B_z_* data.

### Limitations and Future Steps

We used the optimization of model conductivities as a means to estimate the current density distribution, and systematically ensured the quality of the calculated current density distribution by statistical testing. However, this does not imply that the optimized conductivities (or their ratios) are necessarily robust and trustworthy, as conductivity variations of different tissues can have quite similar effects on the current flow patterns and the measured current-induced magnetic fields. For example, *increasing* scalp conductivity and *decreasing* skull conductivity both have the same main effect of increasing the amount of current shunted through the scalp and decreasing the current that enters the intracranial cavity. Alternatively, also *decreasing* the overall conductivity of the intracranial tissues will have a similar impact on the current flow pattern. In other words, various combinations of the tissue conductivities can give similar current flow patterns, rendering a stable estimation of the conductivities in the presence of measurement noise challenging. This limitation does not affect the stability of the estimated current flow distributions. The conductivities are fitted to best explain the magnetic field in the imaged slice that is strongly depending on the current flow in the surrounding region. The simulated current flows in distant regions are thus not necessarily more accurate than simulations based on standard conductivities taken from literature. Similarly, they do not necessarily generalize to other electrode positions, which becomes obvious when comparing the scalp conductivities for the LR and AP injections (Fig. S10). Interestingly, both skull thickness and composition underneath the electrode positions are different for the two injection schemes, which might be reflected in the change of the conductivities.

The main aims of this study were to analyze how well the projected current density algorithm can be applied to in-vivo *B_z_* data from the human intracranial region and to compare it with an approach that relies on the optimization of a low number of parameters (the compartmental tissue conductivities) of anatomically detailed head models. While voxel-wise reconstructions of the current densities or conductivities remain the ultimate aim of MREIT and MRCDI, our findings highlight the importance of tailoring the reconstruction approaches to the quality and specific properties of the measurements. We based our conclusions also on hierarchical statistical testing and in-vivo *B_z_* data, as we feel that this best ensured the validity of our findings. In contrast, for pure simulation studies, it is more challenging to ensure transferability of the results to the envisioned application and to properly account for factors such as deviations of the noise floor from spatially independent and identically distributed Gaussian noise.

Our results suggest that MRCDI data might be very valuable for the further development of personalized volume conductor models of the head. A relevant question to address would be to examine which aspects of the head modeling need to be improved to ensure that the same conductivity values for the modelled tissue compartments are reached independent of electrode montage. Considering that the residuals still contain spatial patterns above the noise floor also for the 4c and 5c head models with optimized conductivities, it seems promising to test the impact of new segmentation methods (Puonti et al., 2020) on data from a larger group of subjects. In addition, as the simulations assume homogenous conductivities of the brain tissues, it might be interesting to explore whether localized changes of the tissue conductivities due to pathologies such as tumors are detectable as outlier regions in the residual images. This effect would be similar to the local increases in the residual images around the ventricles for 3c models and the AP injections seen here. Increased coverage of the imaged region beyond a single slice would be beneficial for these developments. However, this requires a careful tradeoff against maintaining and further increasing the SNR and robustness to physiological noise.

## CREDIT AUTHORSHIP CONTRIBUTION STATEMENT

**Hasan H. Eroğlu:** Conceptualization, Methodology, Software, Validation, Formal analysis, Investigation, Interpretation of results, Writing - Original Draft, Visualization

**Oula Puonti:** Conceptualization, Methodology, Software, Investigation, Interpretation of results, Writing - Review & Editing

**Cihan Göksu:** Software, Investigation, Interpretation of results, Writing - Review & Editing

**Fróði Gregersen:** Software, Investigation, Interpretation of results, Writing - Review & Editing

**Hartwig R. Siebner:** Resources, Interpretation of results, Writing - Review & Editing

**Lars G. Hanson:** Conceptualization, Methodology, Interpretation of results, Investigation, Writing - Review & Editing, Supervision

**Axel Thielscher:** Conceptualization, Methodology, Software, Investigation, Interpretation of results, Writing - Original Draft, Writing - Review & Editing, Supervision, Funding acquisition

## ACKNOWLEDGEMENTS

This study was supported by the Lundbeck foundation (grants R288-2018-236 to CG and R244-2017-196 and R313-2019-622 to AT), and a PhD stipend from the Sino-Danish Center for Education and Research to FG. HRS holds a 5-year professorship in precision medicine at the Faculty of Health Sciences and Medicine, University of Copenhagen which is sponsored by the Lundbeck Foundation (Grant Nr. R186-2015-2138).

## REFERENCES

Chauhan, M., Indahlastari, A., Kasinadhuni, A.K., Schar, M., Mareci, T.H., Sadleir, R.J., 2018. Low-Frequency Conductivity Tensor Imaging of the Human Head In Vivo Using DT-MREIT: First Study. IEEE Trans. Med. Imaging 37, 966–976. https://doi.org/10.1109/TMI.2017.2783348

Codecasa, L., Rienzo, L. Di, Member, Senior, Weise, K., Member, Student, Gross, S., Haueisen, J., Elettronica, D., Bioingegneria, I., 2015. Fast MOR-based Approach to Uncertainty Quantification in Transcranial Magnetic Stimulation. Compumag2015 3, 1–2. https://doi.org/10.1109/TMAG.2015.2475120

Eyüboğlu, B.M., 2006. Magnetic Resonance Current Density Imaging. WILEY-Encyclopedia Biomed. Eng. WILEY-Encyclopedia Biomed. Eng. (Metin Akay, ed.) 4, 2147–53.

Gao, N., Zhu, S.A., He, B., 2006. A new magnetic resonance electrical impedance tomography (MREIT) algorithm: the RSM-MREIT algorithm with applications to estimation of human head conductivity. Phys. Med. Biol. 51, 3067–3083. https://doi.org/10.1088/0031-9155/51/12/005

Ghanem, R., Higdon, D., Owhadi, H., 2017. Handbook of Uncertainty Quantification. Springer International Publishing.

Göksu, C., Scheffler, K., Ehses, P., Hanson, L.G., Thielscher, A., 2018. Sensitivity Analysis of Magnetic Field Measurements for Magnetic Resonance Electrical Impedance Tomography (MREIT). Magn. Reson. Med. 79, 748–760. https://doi.org/10.1002/mrm.26727

Göksu, C., Scheffler, K., Siebner, H.R., Thielscher, A., Hanson, L.G., 2019. The stray magnetic fields in Magnetic Resonance Current Density Imaging (MRCDI). Phys. Medica 59. https://doi.org/10.1016/j.ejmp.2019.02.022

Göksu, Hanson, Siebner, H.R., Ehses, P., Scheffler, K., Thielscher, A., 2018. Human in-vivo brain magnetic resonance current density imaging (MRCDI). Neuroimage 171, 26–39. https://doi.org/10.1016/j.neuroimage.2017.12.075

Holdefer, R.N., Sadleir, R., Russell, M.J., 2006. Predicted current densities in the brain during transcranial electrical stimulation. Clin. Neurophysiol. 117, 1388–1397. https://doi.org/10.1016/j.clinph.2006.02.020

Ider, Y.Z., Birgul, O., Oran, O.F., Arikan, O., Hamamura, M.J., Muftuler, L.T., 2010. Fourier transform magnetic resonance current density imaging (FT-MRCDI) from one component of magnetic flux density. Phys. Med. Biol. 55, 3177–3199. https://doi.org/10.1088/0031-9155/55/11/013

Jeong, W.C., Sajib, S.Z.K., Katoch, N., Kim, H.J., Kwon, O.I., Woo, E.J., 2017. Anisotropic Conductivity Tensor Imaging of In Vivo Canine Brain Using DT-MREIT. IEEE Trans. Med. Imaging 36, 124–131. https://doi.org/10.1109/TMI.2016.2598546

Jeong, W.C., Sajib, S.Z.K., Kim, H.J., Kwon, O.I., 2014. Focused Current Density Imaging Using Internal Electrode in Magnetic Resonance Electrical Impedance Tomography (MREIT). IEEE Trans. Biomed. Eng. 61, 1938–1946.

Kasinadhuni, A.K., Indahlastari, A., Chauhan, M., Schär, M., Mareci, T.H., Sadleir, R.J., 2017. Imaging of current flow in the human head during transcranial electrical therapy. Brain Stimul. 10, 764–772. https://doi.org/10.1016/j.brs.2017.04.125

Kraft, D., 1988. A Software Package for Sequential Quadratic Programming. Wiss. Berichtswesen d. DFVLR.

Kwon, O.I., Lee, B. Il, Nam, H.S., Park, C., 2007. Noise analysis and MR pulse sequence optimization in MREIT using an injected current nonlinear encoding (ICNE) method. Physiol. Meas. 28, 1391–1404. https://doi.org/10.1088/0967-3334/28/11/006

Kwon, O.I., Sajib, S.Z.K., Sersa, I., Oh, T.I., Jeong, W.C., Kim, H.J., Woo, E.J., 2016. Current density imaging during transcranial direct current stimulation using DT-MRI and MREIT: Algorithm development and numerical simulations. IEEE Trans. Biomed. Eng. 63, 168–175. https://doi.org/10.1109/TBME.2015.2448555

Miranda, P.C., Mekonnen, A., Salvador, R., Basser, P.J., 2014. Predicting the electric field distribution in the brain for the treatment of glioblastoma. Phys. Med. Biol. 59, 4137.

Nathan, S.S., Sinha, S.R., Gordon, B., Lesser, R.P., Thakor, N. V., 1993. Determination of current density distributions generated by electrical stimulation of the human cerebral cortex. Electroencephalogr. Clin. Neurophysiol. 86, 183–192. https://doi.org/10.1016/0013-4694(93)90006-H

Nielsen, J.D., Madsen, K.H., Puonti, O., Siebner, H.R., Bauer, C., Madsen, C.G., Saturnino, G.B., Thielscher, A., 2018. Automatic skull segmentation from MR images for realistic volume conductor models of the head: Assessment of the state-of-the-art. Neuroimage 174, 587–598. https://doi.org/10.1016/j.neuroimage.2018.03.001

Oh, S.H., Lee, B. Il, Woo, E.J., Lee, S.Y., Cho, M.H., Kwon, O., Seo, J.K., 2003. Conductivity and current density image reconstruction using harmonic Bz algorithm in magnetic resonance electrical impedance tomography. Phys. Med. Biol. 48, 3101–3116. https://doi.org/10.1088/0031-9155/48/19/001

Park, C., Lee, B. Il, Kwon, O.I., 2007. Analysis of recoverable current from one component of magnetic flux density in MREIT and MRCDI. Phys. Med. Biol. 52, 3001–3013. https://doi.org/10.1088/0031-9155/52/11/005

Puonti, O., Van Leemput, K., Saturnino, G., Siebner, H., Madsen, K., Thielscher, A., 2020. Accurate and robust whole-head segmentation from magnetic resonance images for individualized head modeling. Neuroimage accepted.

Sajib, S.Z.K., Kim, H.J., Kwon, O.I., Woo, E.J., 2012. Regional absolute conductivity reconstruction using projected current density in MREIT. Phys. Med. Biol. 57, 5841–5859 PHYSICS. https://doi.org/10.1088/0031-9155/57/18/5841

Saturnino, G.B., Madsen, K.H., Thielscher, A., 2019a. Electric field simulations for transcranial brain stimulation using FEM: An efficient implementation and error analysis. J. Neural Eng. 16. https://doi.org/10.1088/1741-2552/ab41ba

Saturnino, G.B., Thielscher, A., Madsen, K.H., Knösche, T.R., Weise, K., 2019b. A principled approach to conductivity uncertainty analysis in electric field calculations. Neuroimage 188. https://doi.org/10.1016/j.neuroimage.2018.12.053

Scott, G.C., Joy, M.L.G., Armstrong, R.L., Henkelman, R.M., 1991. Measurement of nonuniform current density by magnetic resonance. IEEE Trans. Med. Imaging 10, 362–374. https://doi.org/10.1109/42.97586

Thielscher, A., Antunes, A., Saturnino, G.B., 2015. Field modeling for transcranial magnetic stimulation: A useful tool to understand the physiological effects of TMS?, in: Proceedings of the Annual International Conference of the IEEE Engineering in Medicine and Biology Society, EMBS. https://doi.org/10.1109/EMBC.2015.7318340

Thielscher, A., Opitz, A., Windhoff, M., 2011. Impact of the gyral geometry on the electric field induced by transcranial magnetic stimulation. Neuroimage 54, 234–243. https://doi.org/10.1016/j.neuroimage.2010.07.061

Wiener, N., 1938. The Homogeneous Chaos. Am. J. Math. 60, 897–936.

Woo, E.J., Seo, J.K., 2008. Magnetic resonance electrical impedance tomography (MREIT) for high-resolution conductivity imaging. Physiol Meas 29, R1–R26.

Yazdanian, H., Saturnino, G.B., Thielscher, A., Knudsen, K., 2020. Fast evaluation of the Biot-Savart integral using FFT for electrical conductivity imaging. J. Comput. Phys. 411, 1–7. https://doi.org/10.1016/j.jcp.2020.109408

