## Supplementary Material for "On the Reconstruction of Magnetic Resonance Current Density Images of the Human Brain: Pitfalls and Perspectives"

#### Supplementary Material A: Figures

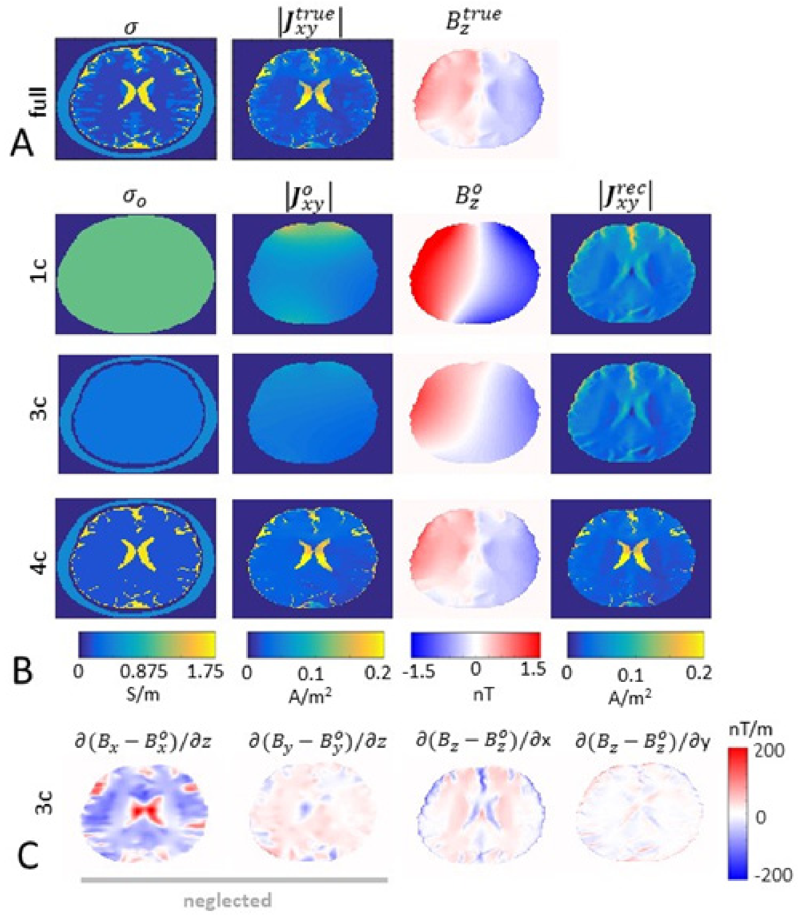


**Figure S1** Current density reconstruction for an AP current injection. (A) Conductivity distribution $\sigma$ of the full model used as ground truth and the corresponding simulated $\left|\boldsymbol{J}_{xy}^{true}\right|$ and $B_z^{true}$ distributions. (B) Results for the 1c, 3c and 4c head models. Columns 1-3 show the conductivity distributions and the corresponding $\left|\boldsymbol{J}_{xy}^{o}\right|$and $B_z^o$ distributions. Columns 4 depicts the reconstructed current densities $|\boldsymbol{J}_{xy}^{rec}|$. (C) Visualization of the terms that are neglected and included, respectively, in the projected current density algorithm (see eq. 3) for the 3c head model.

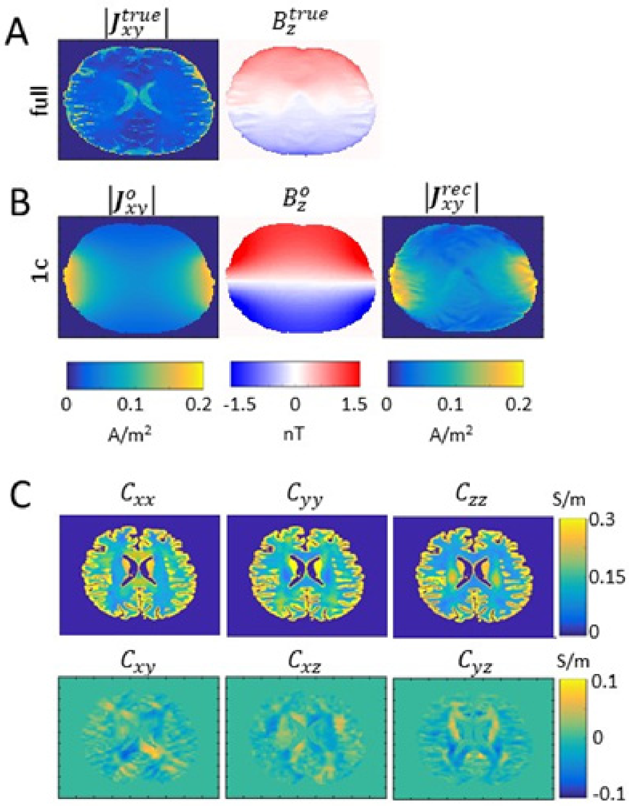


**Figure S2** Results for LR current injection when using a head model as ground truth that incorporated anisotropic brain conductivity derived from diffusion MR data. (A) $|J_{xy}^{true}|$ and $B_z^{true}$ distributions obtained for the full model with anisotropic conductivity distribution of the brain tissues. (B) $|J_{xy}^{o}|$, $B_z^{o}$ and $|J_{xy}^{rec}|$ distributions obtained for the 1c model. $|J_{xy}^{rec}|$ fails to capture the details of $|J_{xy}^{true}|$ ($R^2$=0.06 between $|J_{xy}^{true}|$ and $|J_{xy}^{rec}|$). (C) Conductivity tensors of the brain region of the full model. They were obtained from a diffusion tensor image by means of a volume normalized mapping that keeps the geometric mean of the eigenvalues identical to the standard isotropic conductivity (Güllmar et al., 2010). For the other tissues, the isotropic conductivities listed in the main paper were used. The diffusion images were acquired on a Philips Achieva 3T equipped with a 32 channel head coil and a SE EPI sequence (TR/TE=10s/75ms; flip angle=90°; 70 horizontal slices; phase-encoding direction posterior-anterior; matrix=112x112; voxel size=2x2x2mm$^3$; bandwidth per pixel=23.8 Hz; SENSE=2; 30 volumes with b=1000 s/mm$^2$ and diffusion directions evenly distributed over the "whole sphere"; 4 volumes with b=0 s/mm$^2$). In addition, 3 volumes with b=0 s/mm$^2$ were recorded with opposite phase-encoding direction. Preprocessing was performed using the SimNIBS script *dwi2cond* that calls FSL *topup* and *eddy* to correct static and eddy-current-related distortions (see simnibs.github.io/simnibs/build/html/documentation/command_line/dwi2cond.html for details).

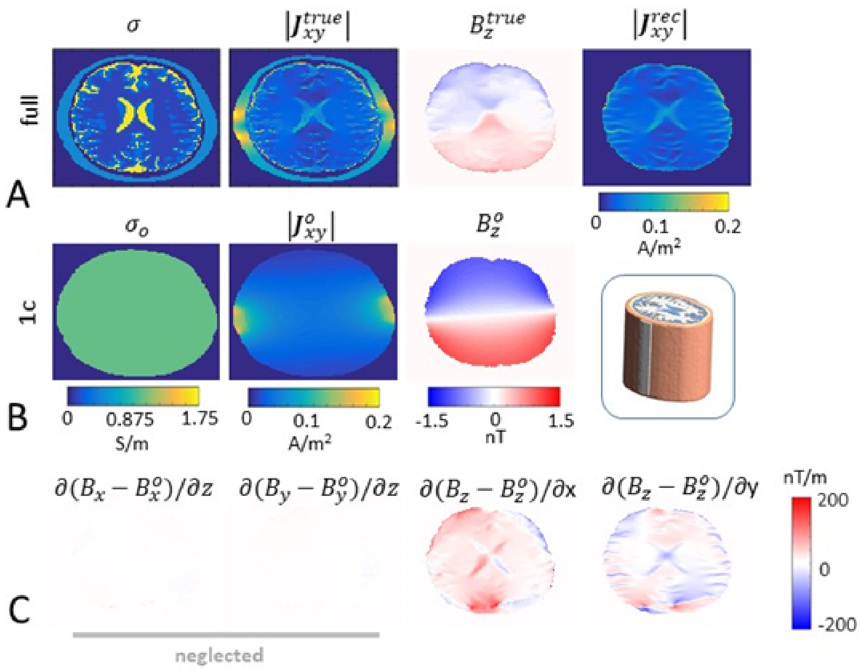


**Figure S3** Current density reconstruction for a simplified "head" model that is uniform along the z-direction. (A) Columns 1-3 show the conductivity distribution for an axial slice of the head model, and the simulated $|J_{xy}^{true}|$ and $B_z^{true}$ for a LR current injection. Column 4 depicts the $|J_{xy}^{rec}|$ distribution reconstructed with the projected current density algorithm ($R^2$=0.63 between $|J_{xy}^{true}|$ and $|J_{xy}^{rec}|$; average strengths: $|J_{xy}^{true}|$ 0.035 A/m², $|J_{xy}^{rec}|$ 0.032 A/m²). (B) Conductivity, current and magnetic fields of the 1c model used for the projected current density algorithm. (C) Visualization of the terms that are neglected and included, respectively, in the projected current density algorithm (see eq. 3) for the 1c head model.

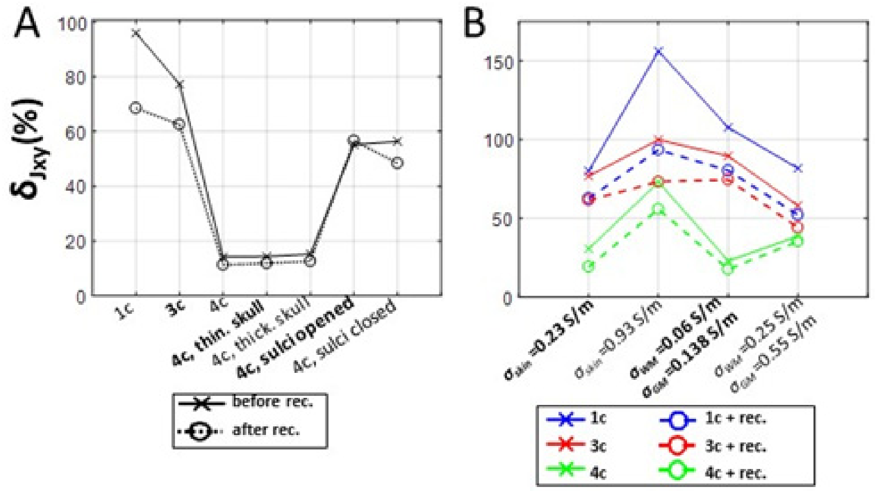


**Figure S4** RMS errors $\delta_{Jxy}$ of different head models for an AP current injection, with and without applying the projected current density algorithm. Solid and dashed lines represent the $\delta_{Jxy}$ values for the $\boldsymbol{J}_{xy}^{o}$ and $\boldsymbol{J}_{xy}^{rec}$ distributions, respectively. (A) Dependence of $\delta_{Jxy}$ on the anatomical level of detail of the head model (1c, 3c and 4c models) and on segmentation errors of the 4c model. (B) Dependence of $\delta_{Jxy}$ on variations of the conductivity of the full model that was used as ground truth.

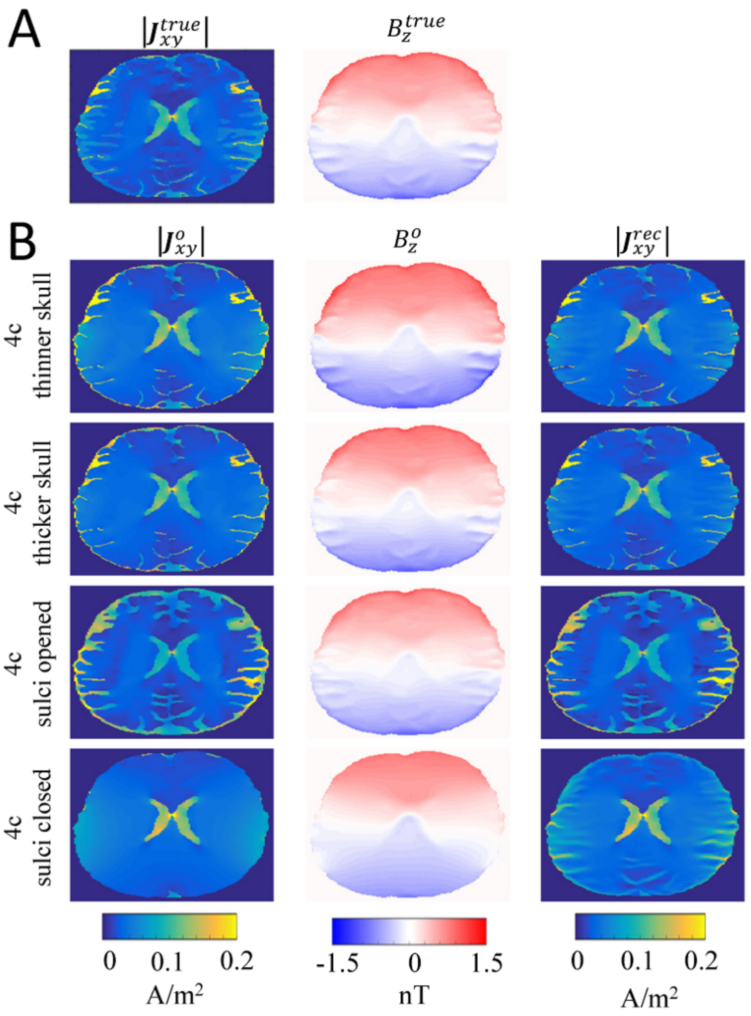


**Figure S5** Current density reconstructions for a LR current injection and head models with segmentation errors. **(A)** $|J_{xy}^{true}|$ and $B_z^{true}$ distributions of the full head model used as ground truth . (B) Distributions of $|J_{xy}^{o}|$, $B_z^{o}$ and $|J_{xy}^{rec}|$ obtained for 4c models with segmentation errors.

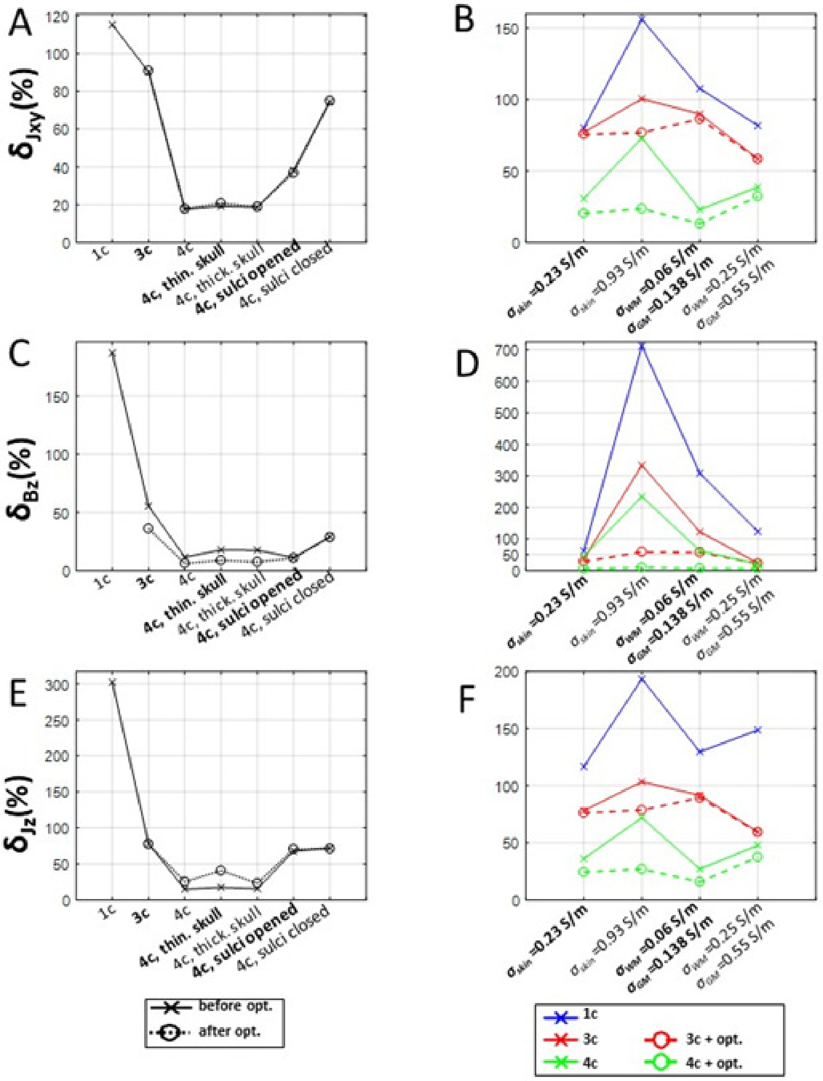


**Figure S6** RMS errors $\delta_{Jxy}$, $\delta_{Bz}$ and $\delta_{Jz}$ of different head models for an AP current injection, with and without optimized conductivities. Solid and dashed lines correspond to the error values obtained for the models with literature and optimized conductivities, respectively. The first column (A, C & E) shows the dependence of $\delta_{Jxy}$, $\delta_{Bz}$ and $\delta_{Jz}$ on the anatomical level of detail of the head model (1c, 3c and 4c models) and on segmentation errors of the 4c model. The second column (B, D & F) shows the dependence of $\delta_{Jxy}$, $\delta_{Bz}$ and $\delta_{Jz}$ on variations of the conductivity of the full model that was used as ground truth.

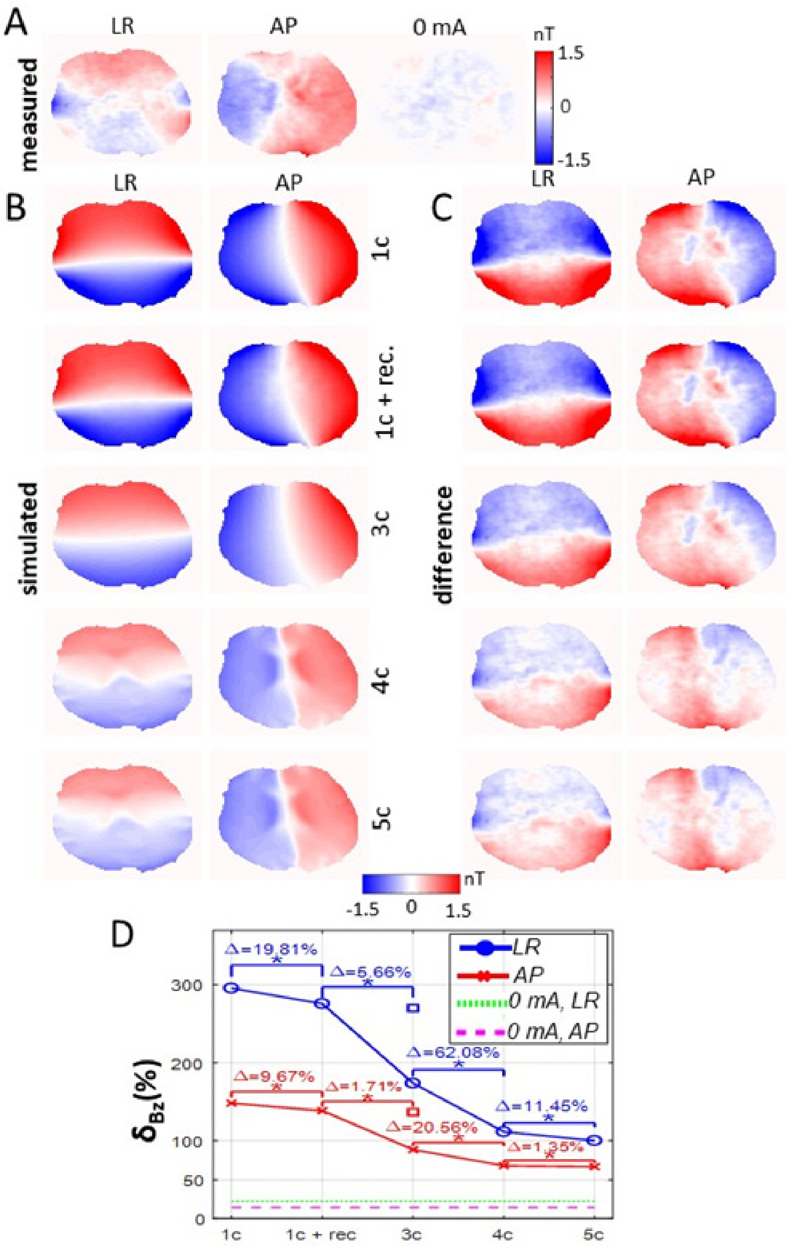


**Figure S7** Experimental $B_z$ measurements and the corresponding simulated $B_z$ distributions for the second subject. The conductivity values of the 3c, 4c and 5c models were chosen from literature values, using the procedure described in Methods section. (A) Measured $B_z$ distributions with and without 1 mA current injections in LR and AP directions. (B) Simulated $B_z$ distributions obtained for the 1c, 3c, 4c and 5c models and for the projected current density reconstruction based on the 1c model (1c+rec.). For the latter, $B_z$ was calculated by using the reconstructed current density $\boldsymbol{J}^{rec}$ in the imaged slice and $\boldsymbol{J}^{o}$ of the 1c head model in the rest of the head volume. (C) Differences between the measured and simulated $B_z$ distributions. (D) Dependence of $\delta_{Bz}$ on the model. Asterisks (*) indicate significant differences between the models at $p \leq$

0.01, Bonferroni corrected for 8 comparisons. The squares (□) represent $\delta_{Bz}$ values obtained for $B_z$ distributions reconstructed using $\boldsymbol{J}^o$ of the 3c model in the imaged slice and $\boldsymbol{J}^o$ of the 1c model in the rest of the head volume. The dashed green and purple lines represent the noise floors $nf$ (eq. 7).

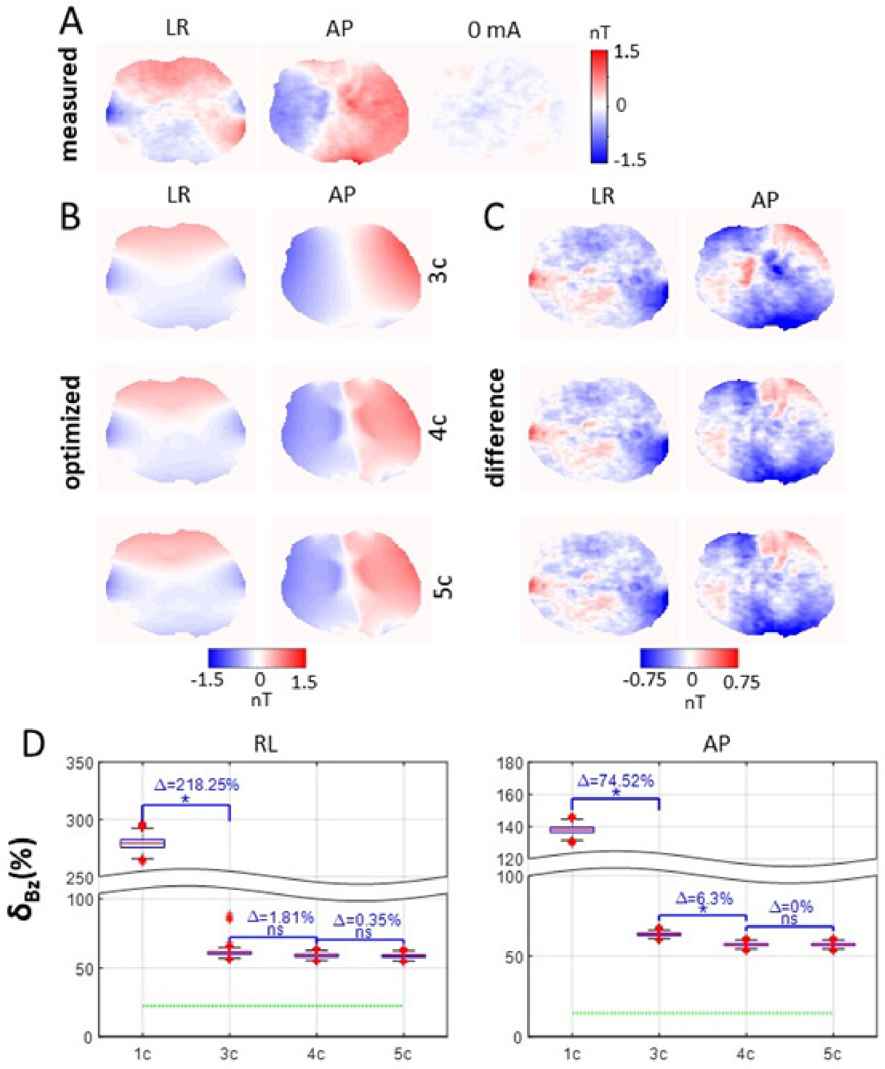


**Figure S8** Results for the 3c, 4c and 5c models with optimized conductivities for the second subject, determined by minimizing the difference to the measured $B_z$ distributions. (A) Measured $B_z$ distributions with and without 1 mA current injections. (B) Optimized $B_z$ distributions for the 3c, 4c and 5c models, corresponding to the median $\delta_{Bz}$ shown in D. (C) Difference between the measured and optimized $B_z$ distributions. (D) Dependence of $\delta_{Bz}$ on the model. The red lines indicate the median, the boxes mark the 25th and 75th percentiles and the whisker lengths is set to 1.5 times the interquartile range. Values outside that range are markeas outliers. Blue asterisks (*) indicate significant differences between the models at $p \leq 0.01$, Bonferroni corrected for 6 comparisons. The dashed green lines indicate the noise floors.

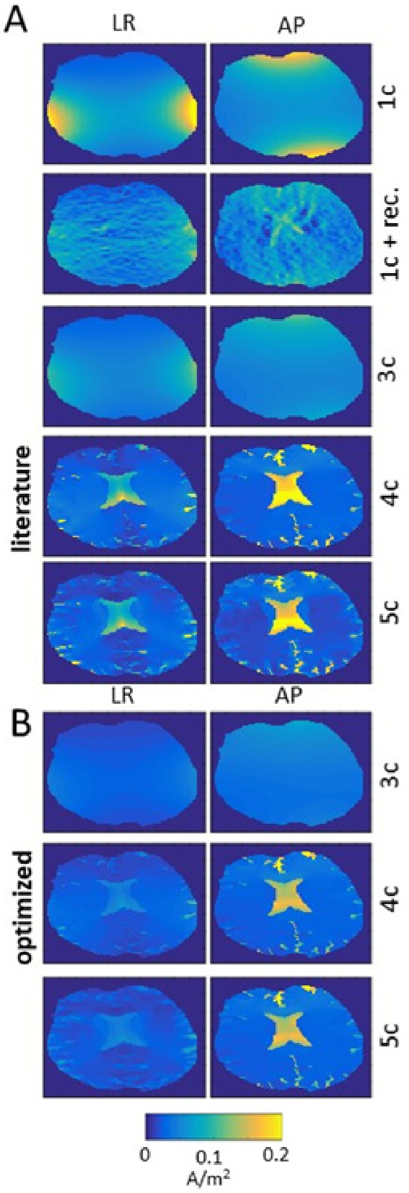


**Figure S9** Estimated current density distributions $|\boldsymbol{J}_{xy}|$ for the second subject. (A) Results for the 1c, 3c, 4c and 5c models based on literature conductivities. Also the results for the projected current density reconstruction based on the 1c model are shown (1c+rec.). The corresponding $B_z$ fields are shown in Fig. S7. (B) Results for the 3c, 4c and 5c models based on optimized conductivities. The corresponding $B_z$ fields are shown in Fig. S8.

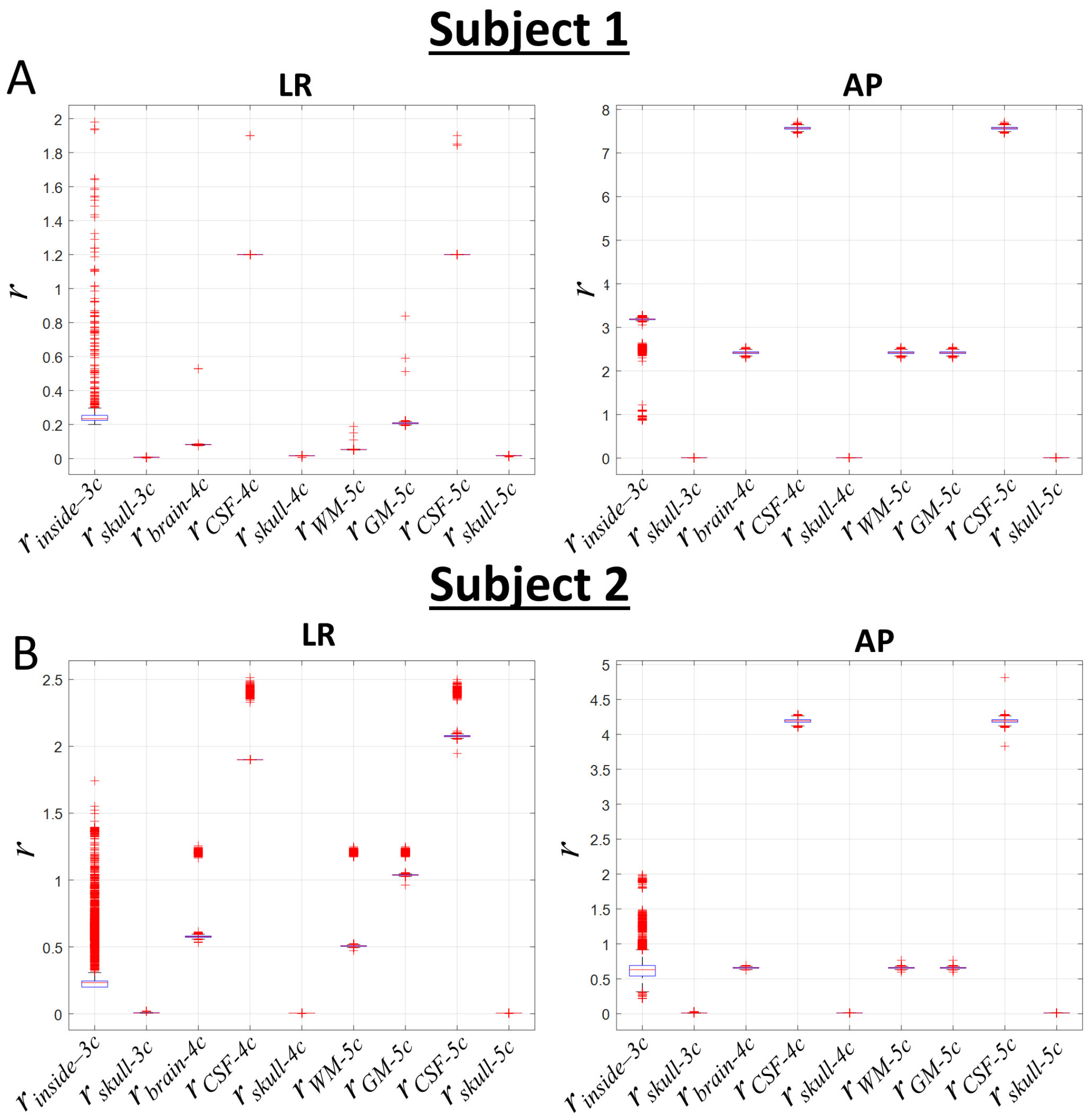


**Figure S10** Optimized conductivity ratios with respect to the optimized scalp conductivities ($r = \frac{\sigma_{compartment}}{\sigma_{scalp}}$) for the 3c, 4c and 5c models of the (A) first and (B) second subjects. Without knowledge of the resistance between the two stimulation electrodes, the tissue conductivities are only uniquely determined up to a common scaling factor. For this reason, they are shown as ratios with respect to the optimized scalp conductivity, which was arbitrarily chosen as reference. The ranges for optimal conductivity ratios differ between LR and AP injections, showing that the resulting values are specific for the tested electrode montage. Such effects might be caused by different skull compositions in the regions under the electrodes that are not accounted for by the head models. The optimized conductivity values should not be taken as accurate estimates of the true biological tissue conductivities. Rather, they merely give the best current flow pattern to approximate the measured $B_z$ field for the specific head model and electrode montage.

### Supplementary Material B: Iterative Projected Current Density Reconstruction Algorithm

Kwon and colleagues (2016) proposed an iterative version of the projected current density algorithm that was aimed at estimating the current flow in the human brain during transcranial electric stimulation (TES). It is here demonstrated that the algorithm requires $B_z^m$ measurements in the complete head volume in order to provide useful results. The algorithm starts with an initial estimation of the current flow $\boldsymbol{J}^0$ using an anatomically detailed head model. This current flow estimate is then updated by repeatedly applying the projected current density algorithm until the relative difference between the current flow estimates of two successive iterations reaches a pre-defined threshold. The first iteration is identical to applying the standard projected current density algorithm:

$$\boldsymbol{J}^1 = \boldsymbol{J}^o + \frac{1}{\mu_o}\left[\frac{\partial(B_z^m - B_z^o)}{\partial y}, \frac{-\partial(B_z^m - B_z^o)}{\partial x}, 0\right] \quad \text{(S.1)}$$

By applying the Biot-Savart law to the updated current density estimate $\boldsymbol{J}^1$, a new estimate of the magnetic flux density $B_z^1$ is obtained. Both $\boldsymbol{J}^1$ and $B_z^1$ are then again used to get a newly updated $\boldsymbol{J}^2$, etc.:

$$\boldsymbol{J}^{n+1} = \boldsymbol{J}^n + \frac{1}{\mu_o}\left[\frac{\partial(B_z^m - B_z^n)}{\partial y}, \frac{-\partial(B_z^m - B_z^n)}{\partial x}, 0\right], \quad \text{(S.2)}$$

where $\boldsymbol{J}^n$ and $B_z^n$ are the projected current density and the corresponding magnetic flux density calculated in the $n^{th}$ iteration. The algorithm stops when the relative difference between once $\boldsymbol{J}^{n+1}$ and $\boldsymbol{J}^n$ converges below a pre-defined tolerance $\frac{|J^{n+1} - J^n|}{|J^n|} < \epsilon$. In the original paper, the algorithm was validated using numerical simulations. It was shown that the solution converged close to the known ground truth ($\boldsymbol{J}^{true}$ and $B_z^{true}$) also in the presence of zero-mean Gaussian noise in the "measured" magnetic flux density $B_z^m$ and when the initial estimation of the current flow $\boldsymbol{J}^0$ was inaccurate due to on purpose incorrectly chosen tissue conductivities.

Our aim here was to test the performance of the algorithm for more realistic $B_z^m$ data that covers only the intracranial compartment (i.e. lacking information about the magnetic flux in the skin and skull) or stems from a single measurement slice. We started by simulating a LR current injection for the full head model shown in Figure 1A as ground truth $\boldsymbol{J}^{true}$ and $B_z^{true}$. Then, we applied the iterative projected current density algorithm to the 4 compartment (4c) head model for evaluation, providing $B_z^m$ data (i) either in the whole volume of the head model (skin + skull + CSF + GM + WM), (ii) only for positions in the intracranial compartment (CSF+ GM + WM), or (iii) only for positions in the intracranial compartment in an axial slice at the level of the electrodes. For the second and third cases, we were able to apply the projected current density algorithm only to update the current flow in a limited volume. Therefore, we determined $\boldsymbol{J}^{n+1}$ as

$$\boldsymbol{J}^{n+1} = \begin{cases} \boldsymbol{J}^{n} + \frac{1}{\mu_o}\left[\frac{\partial(B_z^m - B_z^n)}{\partial y}, \frac{-\partial(B_z^m - B_z^n)}{\partial x}, 0\right] \; \ldots inside\ the\ measurment\ volume \\ \boldsymbol{J}^{o} \; \ldots otherwise \end{cases} \quad (S.3)$$

and then calculated $B_z^{n+1}$ by applying Biot-Savart to $\boldsymbol{J}^{n+1}$. We terminated the algorithm using a stopping criterion of $\epsilon$ = 1 %. In order to evaluate the performance of the iterative algorithm, we calculated the errors

$$\Delta_{Jxy} = \sqrt{\frac{\left[\left(J_x - J_x^{true}\right)^2 + \left(J_y - J_y^{true}\right)^2\right]}{J_x^{true^2} + J_y^{true^2}}} \times 100\ \% \quad (S.4)$$

and

$$\Delta_{B_z} = \sqrt{\frac{\left(B_z - B_z^{true}\right)^2}{B_z^{true^2}}} \times 100\ \% \quad (S.5)$$

along with their corresponding spatial averages, $\delta_{Jxy}$ and $\delta_{Bz}$ (see eqn. 4&6) in the brain region (WM+GM+CSF) of the slice being imaged.

In initial tests, we started the algorithm with $\boldsymbol{J}^0$ data calculated with a 4c head model that was matched as close as possible to the full model by setting the conductivity of the "brain" compartment to the volume or slice average of the WM and GM compartments of the full model and using the same conductivities otherwise. The algorithm converged close to the ground truth after few iterations for all three cases (full head, intracranial volume, axial slice), demonstrating its correct implementation.

We then went on and initialized the algorithm with scalp and skull conductivities that were 20% higher and 20% lower than those of the full model used as ground truth. For each of the three cases, the algorithm started with the same $\left|\boldsymbol{J}_{xy}^1\right|$ (see the first columns of Figure S11 B&C and the first row of Figure S12) in the brain region (WM + GM + CSF), as expected. RMS errors were $\delta J_{xy}^1 = 10.6$ for all three cases (see the first row of Figure S12). For calculation of $B_z^1$ in the second and third cases, the erroneous $\boldsymbol{J}^o$ distributions had to be used (eq. S3) as estimates of the unknown true current density in skin and skull segments.

Correspondingly, also the differences between of $B_z^1$ and the "measured" $B_z^{true}$, indexed by the $\Delta_{B_z^1}$ distributions (second column of Fig. S11C), showed much larger spatial variations for these cases compared to the first case. Expectedly, the RMS errors were also increased ($\delta B_z^1 = 11.35\ \%, 29.51\ \%, 31.6\ \%$ for the first, second, and third cases; see the second row of Fig. S12). With an increasing number of iterations, the algorithm expectedly improves the $\Delta_{B_z}$ distributions in all three cases. However, it improves $\Delta_{Jxy}$ only for the case with complete head volume coverage, but increases it from the second iteration on for the cases with limited volume coverage (third column of Fig. S11 B&C and first row of Fig. S12).

For the case with complete coverage, the algorithm can update the $\boldsymbol{J}^n$ distributions everywhere (including skin and skull) which also globally reduces the difference between the simulated and measured magnetic fields and ensures correct convergence. On the other hand, when $B_z^m$ measurements are only available from a limited volume, the algorithm compensates for the impact of the incorrectly estimated current distributions ($\boldsymbol{J}^o$) outside of that volume on the local magnetic field in order to reduce the difference between the spatial gradients of the simulated and measured fields (eq. S.3). As result, it converges to an incorrect current density distribution.

To summarize, the iterative projected current density reconstruction algorithm requires $B_z^m$ measurements in the complete head volume in order to ensure correct convergence. In contrast, for a limited volume coverage, even moderate deviations of the initial scalp and skin conductivities by 20% from the ground truth as tested here resulted in convergence to an incorrect current density distribution.

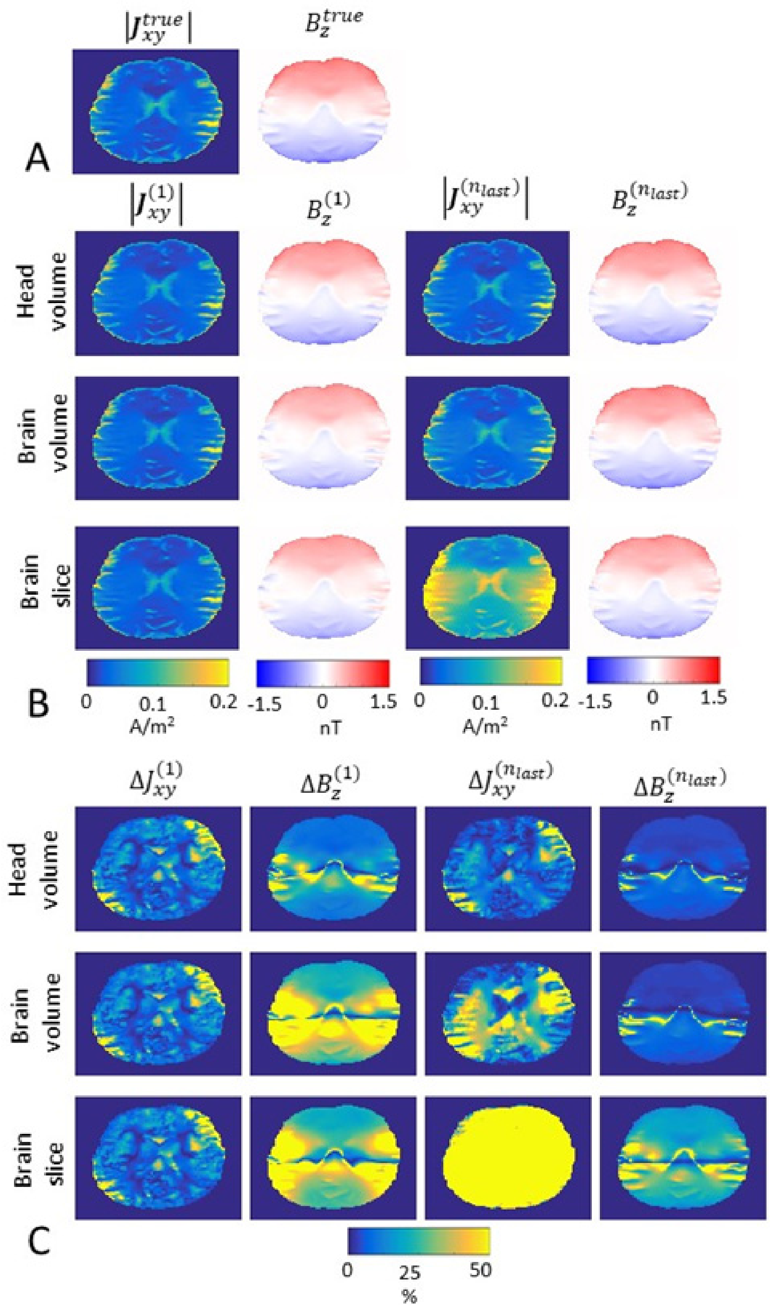


**Figure S11** Current and magnetic flux density reconstructions and corresponding errors for the iterative reconstruction algorithm. (A) $\left|J_{xy}^{true}\right|$ and $B_z^{true}$ distributions for a LR current injection in the full head model used as ground truth. (B) Distributions of the reconstructed current and magnetic flux densities for the first ($\left|J_{xy}^{(1)}\right|$, $B_z^{(1)}$) and last iterations ($\left|J_{xy}^{(n_{last})}\right|$, $B_z^{(n_{last})}$). (C) Error distributions of the reconstructed current and magnetic flux densities for the first ($\Delta J_{xy}^{(1)}$, $\Delta B_z^{(1)}$) and last iterations ($\Delta J_{xy}^{(n_{last})}$, $\Delta B_z^{(n_{last})}$).

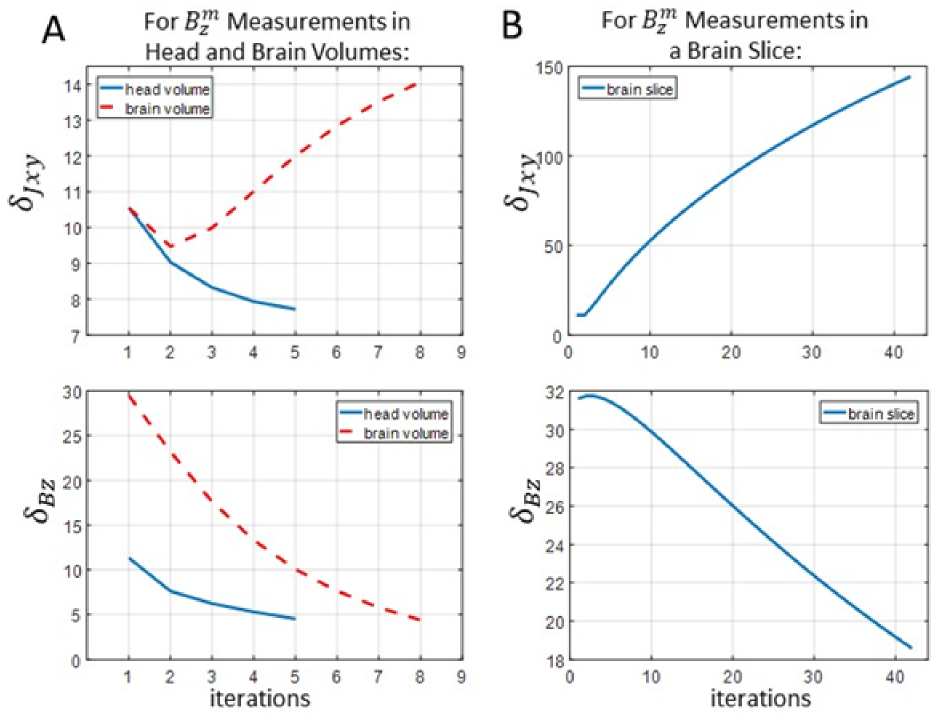


**Figure S12** RMS errors $\delta_{Jxy}$ and $\delta_{Bz}$ for the iterative reconstruction algorithm. In column (A), $B_z^m$ measurements are available in the whole head (skin + skull + CSF + GM + WM) or the whole brain volume (CSF+ GM + WM), whereas in column (B) $B_z^m$ measurements are only available in an axial slice of the brain volume. The algorithm converged to $\epsilon = 1$ % at 5, 8, and 42 iterations for the first (head volume coverage), second (brain volume coverage) and third (brain slice) cases, respectively.

#### References

Güllmar, D., Haueisen, J., Reichenbach, J.R., 2010. Influence of anisotropic electrical conductivity in white matter tissue on the EEG/MEG forward and inverse solution. A high-resolution whole head simulation study. Neuroimage. https://doi.org/10.1016/j.neuroimage.2010.02.014

Kwon, O.I., Sajib, S.Z.K., Sersa, I., Oh, T.I., Jeong, W.C., Kim, H.J., Woo, E.J., 2016. Current density imaging during transcranial direct current stimulation using DT-MRI and MREIT: Algorithm development and numerical simulations. IEEE Trans. Biomed. Eng. 63, 168–175. https://doi.org/10.1109/TBME.2015.2448555
